# Age-associated increases in inter-individual gene expression variability across human tissues

**DOI:** 10.64898/2026.04.14.718522

**Authors:** Josh Bartz, Paola Rivera, Laura J. Niedernhofer, Lei Zhang, Xiao Dong

**Affiliations:** Masonic Institute on the Biology of Aging and Metabolism, University of Minnesota, Twin Cities, Minneapolis, MN 55455, USA; Department of Genetics, Cell Biology and Development, University of Minnesota, Twin Cities, Minneapolis, MN 55455, USA; Department of Biochemistry, Molecular Biology, and Biophysics, University of Minnesota Twin Cities, Minneapolis, MN 55455, USA

## Abstract

Aging involves progressive physiological decline, yet the underlying transcriptomic patterns remain poorly understood. Although differentially expressed genes (DEGs) have been the primary focus of previous studies, here we investigate differentially variable genes (DVGs) using a novel Gene Stability Score (GSS). In 30 tissue types from nearly 1,000 individuals in the Genotype-Tissue Expression (GTEx) project, age- and sex-related DVGs account for approximately 15% of overall expression variability between samples of the same tissue, with age-related DVGs specifically contributing 7.7%. We further show that DEGs and DVGs affect distinct biological pathways, and that inter-individual instability is significantly correlated with cell-to-cell transcriptional noise. Moreover, gene regulatory network analysis reveals that this variability is not random but is shaped by local network architecture. Finally, we identify robust reference genes, including *TBP, PUM1*, and *TMEM199*, for RT-qPCR experiments in studying age-related gene expression changes in humans. Together, our findings suggest that aging involves both coordinated transcriptional programs and increased stochasticity across individuals and cells.

## MAIN

Aging is a complex biological process characterized by progressive decline of a wide range of critical physiological functions^1–3^. A set of age-related hallmarks that capture key changes underlying the aging process has been proposed^4^. Among these, genomic and epigenomic instability represent fundamental molecular-level alterations; however, the mechanisms by which they drive the aging process remain unclear^5,6^. The effects of these stochastic alterations likely converge on the disruption of gene transcriptional regulation^5,7^. Yet, the exact patterns of age-related transcriptomic alterations remain poorly understood^8^.

Previous investigations of the aging transcriptome have overwhelmingly focused on identifying age-associated differentially expressed genes (DEGs). While valuable, this approach overlooks genes whose expression variability changes with age, referred to as differentially variable genes (DVGs), in contrast to stable genes (SGs). These genes likely represent more direct consequences of genomic and epigenomic instability, particularly at the single-cell level^9–15^. However, only a limited number of studies have examined transcriptomic instability. Moreover, findings from single-cell analyses remain inconsistent: some studies report increased transcriptional variability with age, whereas others do not^16–21^. This discrepancy is likely due to the high level of technical noise inherent in single-cell transcriptomic measurements, which can obscure true biological variability. Importantly, evidence for age-associated transcriptomic instability has also been observed at the bulk RNA level. For example, a landmark study of monozygotic twins showed that gene expression profiles diverge substantially with age, with older twin pairs exhibiting markedly greater transcriptional differences than younger pairs despite sharing identical genomes^22^.

However, it remains unclear which genes exhibit increasing inter-individual transcriptional variability with age within the same tissue, and whether these genes are related to those showing increased instability at the single-cell level. To address this question, we developed a stability metric, Gene Stability Score (GSS), to quantify gene expression variability within a human population independent of age information. Currently, no widely accepted method exists to quantify gene expression stability in bulk RNA-sequencing datasets, so we designed this metric by leveraging algorithms originally developed to discover reference genes used in Reverse Transcription-quantitative Polymerase Chain Reaction (RT-qPCR) experiments. Applying this metric to the Genotype-Tissue Expression (GTEx) dataset, we show that inter-individual transcriptional variability within the same tissues across the GTEx cohort arises from both age- and sex-associated DEGs and age- and sex-associated DVGs (**Figure 1A**). Furthermore, by analyzing published single-cell datasets, we demonstrate a significant correlation between these age-related inter-individual DVGs and age-related cell-to-cell DVGs.

**Figure 1.**
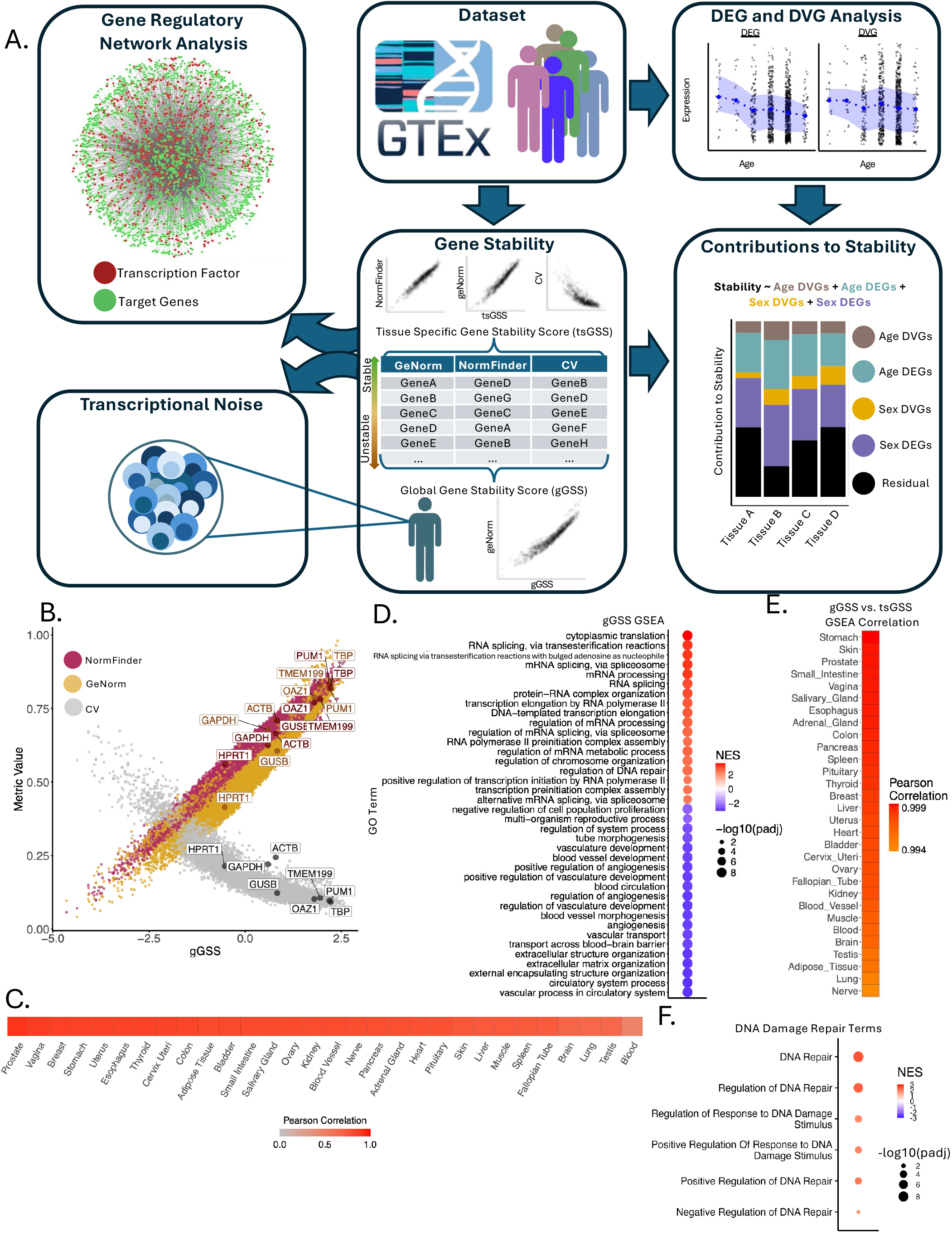
Gene expression stability is consistent across tissues. (A) Schematic overview of the study. (B) gGSS correlates strongly with its underlying stability metrics. (C) tsGSS for each tissues has a strong correlation with gGSS, suggesting that much of expression stability is conserved between tissues and that the gGSS is a good proxy for general gene stability. (D) Dotplots of GO terms enriched for genes ranked by their gGSS, highlighting 38 GO terms that were enriched in all 31 tsGSS GSEA analyses. Positive enrichment represents pathways overrepresented in stable genes while negative enrichment represents pathways overrepresented in unstable genes. (E) Correlation between pathway enrichment for the 38 selected terms derived from gGSS and tsGSS. For each tissue, gene set enrichment analysis was performed using tsGSS values across the same GO terms shown in (D). Normalized enrichment scores (NES) from each tissue-specific analysis were then compared to the NES obtained from the global gGSS-based enrichment. The resulting correlations quantify the concordance between global and tissue-specific stability-associated pathway enrichment, with higher correlations indicating greater agreement in pathway-level signals across tissues. (F) Enrichment of DNA damage repair related pathways based on gGSS.

## RESULTS

### Measuring Gene Stability

To quantify transcriptional variability of a gene across a population, we developed a single measurement that integrates three variability/stability measurements: GeNorm^23^, Normfinder^24^, and the coefficient of variation (CV). GeNorm and Normfinder were developed to identify transcriptionally stable genes, which can be used as references in RT-qPCR experiments. We limited the following analyses to the 9,000 genes with the highest average expression across GTEx, avoiding quantification noise from lowly expressed genes and tissue-specific expression.

We applied the three methods to each tissue independently, across all tissue samples regardless of sex or age, allowing us to assess gene stability across both variables (sex and age). While highly correlated, they do not give identical results (**Supplementary Figure 1**). To then capture a single unified measure of stability, we performed principal component analysis (PCA) and the first principal component (PC1) explained the vast majority of variation shared across methods. (PC1; **Supplementary Figure 2**). Therefore, PC1 captures the shared signal across methods and represents a consensus of the three measurements. We used this PC1 as a unified measurement of gene variability for each tissue, denoted as the tissue-specific Gene Stability Score (tsGSS). A high GSS indicates that a gene maintains stable expression across human subjects, whereas a low GSS reflects greater variability, corresponding to unstable gene expression. Across the 30 analyzed tissues, tsGSS exhibits a high degree of consistency (**Supplementary Table 1**), indicating that genes stable in one tissue are more likely to be stable in others. Given the observed consistency in gene stability across tissues, we performed another PCA reduction on the GeNorm, Normfinder, and CV of the 30 tissue types. Over 50% of the variation was captured in the PC1 alone (**Supplementary Figure 3**), and subsequently, used as the global Gene Stability Score (gGSS; **Figure 1B**). The gGSS has strong to moderate correlation with all tsGSS, indicating that gGSS does not mask any outlier tissues (**Figure 1C; Supplementary Figure 4**). As an additional validation step, we compared gGSS values derived from the GTEx dataset with expression stability measurements calculated using geNorm, NormFinder, and CV from the Human Protein Atlas^25^ and observed a high degree of similarity (average rho: 0.634, p-value: < 1×10^-10^; **Supplementary Figure 5)**. This suggests that the majority of the calculated expression stability reflects genuine underlying biological variation rather than technical artifacts of the GTEx dataset.

Notably, the most stable gene identified was *WDR20*, which functions to regulate the USP12-UAF1 deubiquitinating enzyme complex^26^, and is important for stabilizing c-Myc and preventing cellular senescence^27^. The second and third most stable genes were *SNW1* and *MED6*, both of which play critical roles in RNA transcription (**Supplementary Table 2**). Additionally, we evaluated the gGSS of commonly used reference genes, including *ACTB, GAPDH, GUSB, HPRT1, OAZ1, PUM1, TMEM199*, for RT-qPCR to assess their suitability as references in age-related studies: they are used as references in typical RT-qPCR experiments under the assumption that they are stably expressed across tissues and age, but this assumption may not be accurate because their transcription stability has been validated mostly in young tissues in addition to a few senescence models^28,29^. Our analysis demonstrates that across all tissues, *TBP, Pum1*, and *TMEM199* are the most appropriate internal references for measuring relative expression changes with age (**Figure 1B**). We also report tissue-specific stability results to guide reference gene selection in individual tissues (**Table 1; Supplementary Table 1**).

**Table 1.**
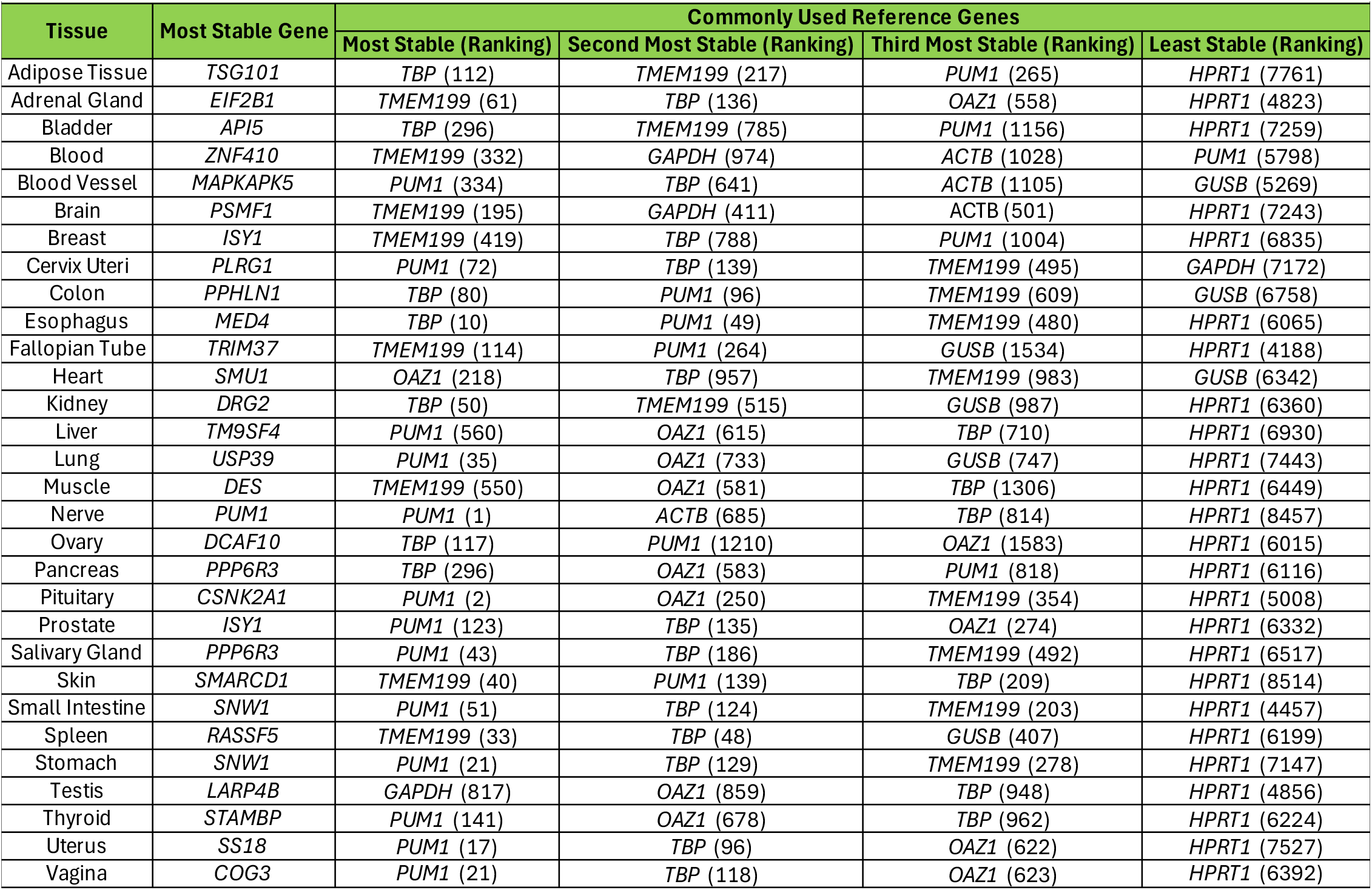
Select stable and unstable genes in each tissue.

In addition, we performed gene set enrichment analysis (GSEA) of genes based on their gGSSs. Stable genes were significantly enriched for multiple RNA processing and splicing pathways, while unstable genes showed strong enrichment for pathways related to blood vessel and vascular development **(Figure 1D)**. GSEA with tsGSS shows a highly similar pattern as the gGSSs, suggesting that the observed pathway enrichments are primarily driven by inter-individual differences rather than tissue-specific effects **(Figure 1E)**. Of note, stable genes are also enriched for multiple DNA damage repair pathways, suggesting this critical process is tightly regulated even during the aging process **(Figure 1F)**.

### DEG and DVGs across in Sexes and Ageing Contribute to Gene Stability

To disentangle the contribution of age- and sex-related DEGs and DVGs to gGSS, we calculated them in the same GTEx dataset using DESeq2^30^ to identify DEGs and the standard deviation to identify DVGs (**Methods**). We then employed a multiple linear regression framework, which assesses the relative importance of predictors in explaining variance in the gGSS (**Figure 2A; Methods**). This analysis revealed that although most gene expression alterations are driven by DEGs (changes in mean expression), DVGs (changes in expression variability) also play a substantial role, accounting for approximately 15% of all expression variation between individuals within the same tissue. Notably, age-related DVGs alone represented 7.7% of all expression variation among the GTEx population (**Figure 2B**). The contribution of DVGs to the tsGSS are similar to their contributions to gGSSs across tissues. At the pathway level, the relative contributions of DEGs and DVGs to most individual pathways closely mirrored the results observed at the whole-tissue level (**Figure 2C**). Of note, the same genes can be both age-related DEGs and DVGs, and this overlap is higher than expected by chance alone (**Supplementary Table 3**). However, most of them are not overlapping and show different distributions in pathways as shown below.

**Figure 2.**
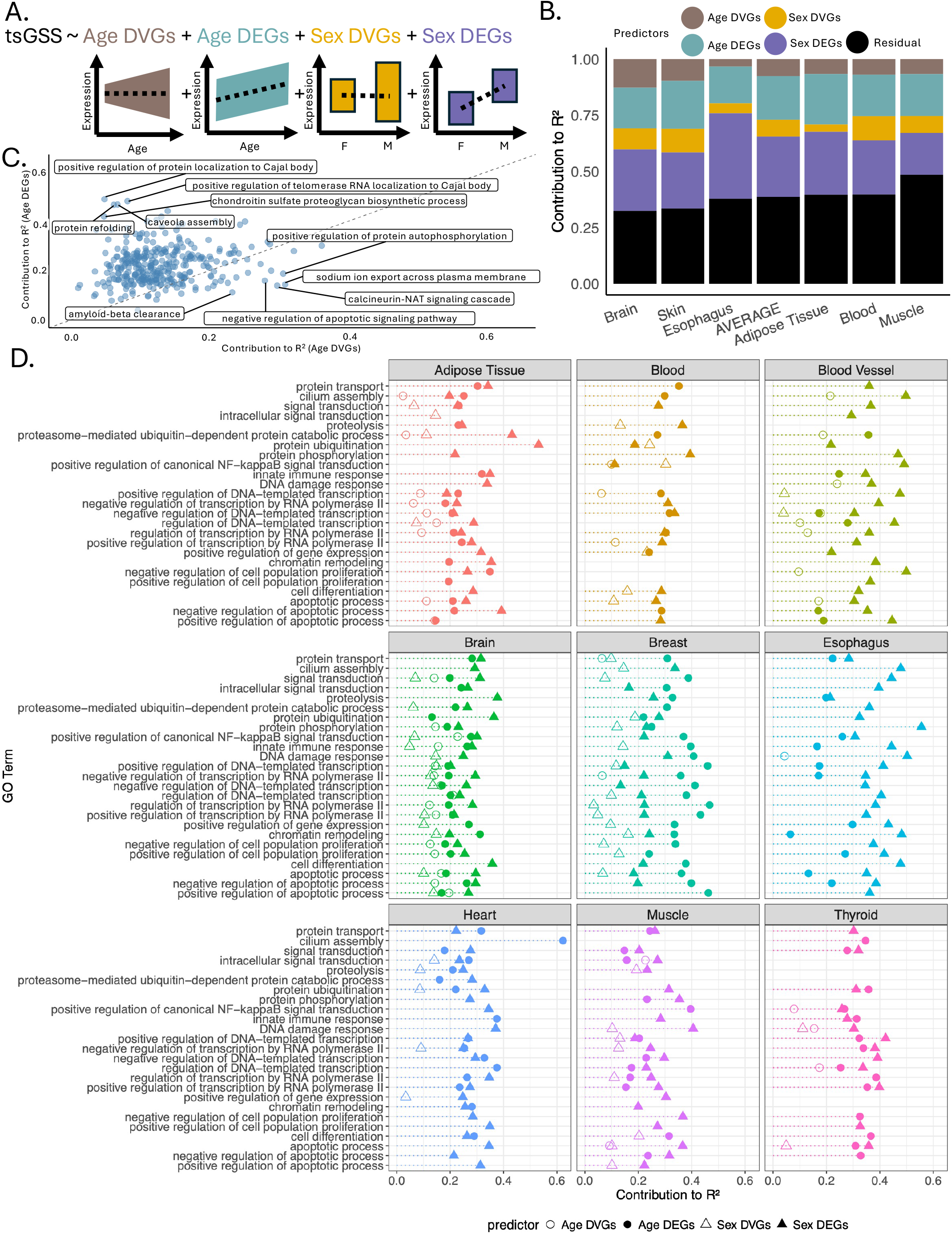
Contributions of DEGs and DVGs to expression variation in GTEX. (A) Schematic overview of multiple linear regression framework. (B) Contributions of age- and sex-associated DEGs and DVGs to total expression variance within individual tissues in the GTEx dataset. Only tissues for which all four predictors (age DEGs, age DVGs, sex DEGs, and sex DVGs) showed statistically significant associations (padj < 0.05) with gGSS in the regression model are displayed. (C) Contributions of age- and sex-associated DEGs and DVGs in explaining expression variance across GO terms within individual tissues. Displayed GO terms are restricted to those with at least 20 statistically significant associations with age- or sex-associated DEGs or DVGs across tissues. Only tissues with at least 700 genes meeting significance and quality criteria for inclusion in the multiple linear regression are displayed. (D) Contributions of age-related DEGs and DVGs to expression variance within GO terms aggregated across all tissues.

We compared the relative contribution of age-related DEGs and DVGs at the pathway level across all tissue types. Pathways dominated by DEGs were primarily associated with core cellular maintenance processes, including proteostasis, cellular organization, and RNA processing. These pathways likely reflect coordinated, directional shifts in fundamental cellular maintenance systems that occur broadly during aging. In contrast, pathways dominated by DVGs include the calcineurin-NFAT signaling cascade, sodium ion export across plasma membrane, amyloid-beta clearance, and negative regulation of apoptotic signaling. (**Figure 2D**). These pathways showed increased inter-individual variability with age, suggesting that processes associated with signaling and homeostasis may become more variable over time. Taken together, these results suggest that aging may involve two complementary modes: a shared, coordinated transcriptional program reflected in DEGs, and a more heterogeneous component reflected in DVGs, potentially arising from stochastic events at the genome level such as DNA damage.

### The Gene Regulatory Network Influences Gene Stability

The gene regulatory network (GRN) has been hypothesized to play a role in gene expression stability^5,12^. To test this, we used the TRRUST database, which contains 8,015 interactions between 2,723 genes^31^. These interactions were obtained using a sentence-based text mining algorithm that collected the interaction between transcription factors (TFs) and target genes (TGs) from the literature.

Using this database, we sought to uncover if genes of similar stability collocated within the GRN. To evaluate whether a gene’s local GRN consists of genes with similar stability profiles, we developed a stability similarity metric (**Methods**). This metric quantifies the degree to which the stability of a given gene aligns with that of its network neighbors. A positive stability similarity score indicates that a gene is more similar in stability to its local network than would be expected by random chance, whereas a negative score suggests the gene’s stability is more divergent from its local GRN. We applied the stability similarity score across all tissues to investigate the role of GRNs in shaping DEGs and DVGs. Overall, DEGs tend to be more dissimilar in stability from their local GRN neighborhoods, whereas DVGs more frequently exhibit stability profiles similar to neighboring genes (**Figures 3A and 3B; Supplementary Table 4**). This suggests that variability is shaped by local GRN organization, possibly due to increased susceptibility of certain regulatory motifs or genes to damage. This damage may then elevate expression variability, which can then propagate through the local network. Conversely, differential expression more often reflects gene-specific regulatory responses that are less coordinated with the surrounding network. Representative results from blood, colon, and skin are shown in **Figures 3C-E**, while results for all tissues are provided in **Supplementary Figure 6**.

**Figure 3.**
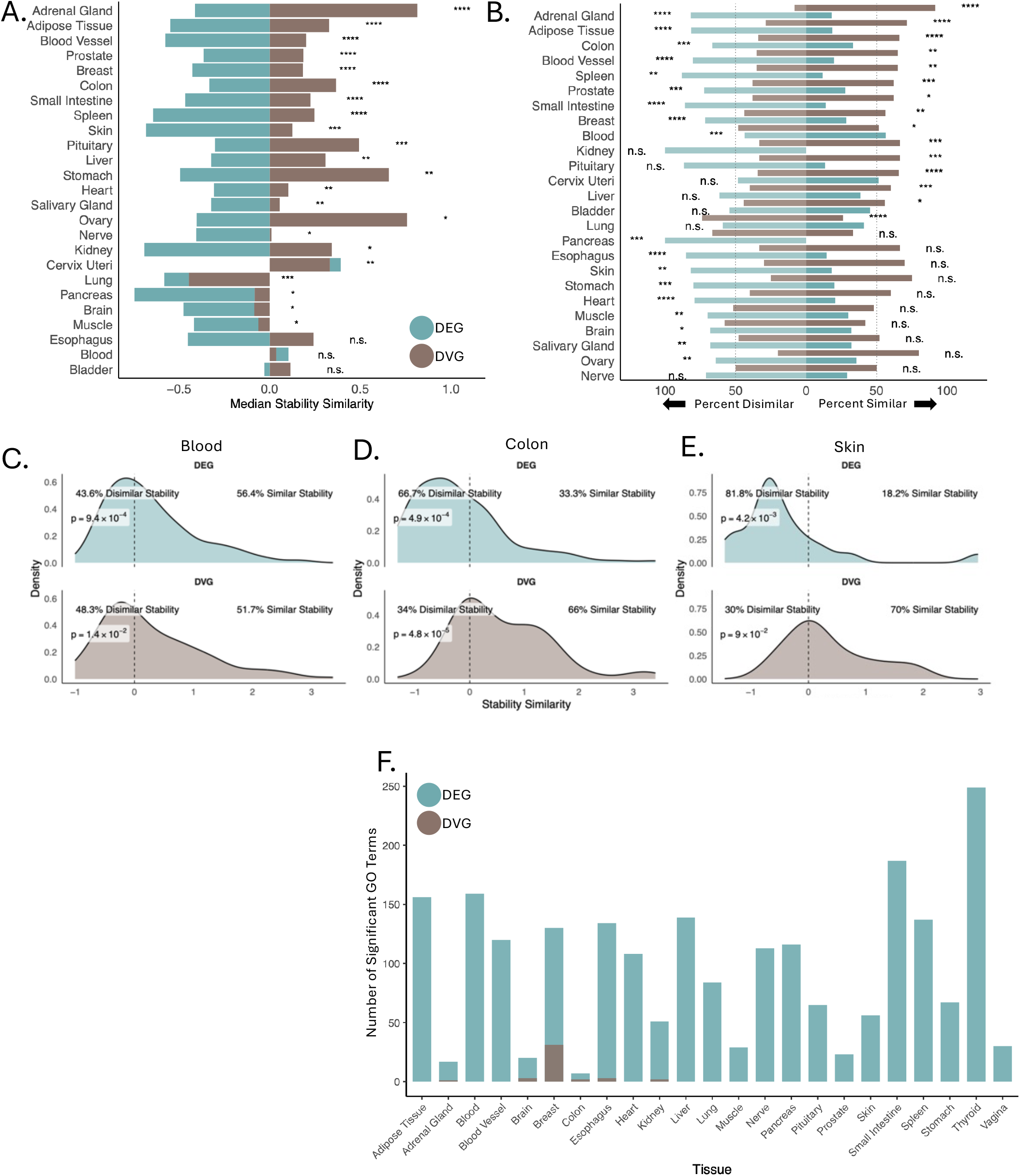
Correlation between gGSS and local regulatory networks. (A) Comparison of stability similarity for DEGs and DVGs within a tissue. Wilcox tests were used to quantify differences in the distributions between DEGs and DVGs (*p < 0.05, **p < 0.01, ***p < 0.001, ****p< 0.0001). (B) Directional bias of stability similarity within DEGs and DVGs. For each group, the proportion of genes with positive and negative stability similarity scores is shown. To determine whether stability scores are systematically shifted toward positive or negative values, we evaluated deviation from a zero-centered distribution using a one-sample Wilcoxon test (*p < 0.05, **p < 0.01, ***p < 0.001, ****p< 0.0001). Density plots of stability similarity for (C) blood, (D) colon, and (E) skin. Displayed p-values are from a two-sided Wilcox test assessing whether the observed stability similarity distribution differs from 0. (F) Number of GO terms significantly enriched in the top 100 DEGs and top 100 DVGs in each tissue.

Importantly, we tested if DEGs and DVGs have different enrichment levels in biological pathways. To avoid any bias due to the difference in their gene numbers, we selected the top 50 most strongly upregulated and top 50 most strongly downregulated DEGs, and the top 50 most increasingly and decreasingly variable DVGs. DEGs show strong enrichment in known biological pathways, consistent with coordinated changes in aging-related programs (median of 108 statistically significantly enriched pathways across tissues). In contrast, DVGs show limited enrichment in pathways (median of 0), suggesting that they may arise not from coordinated shifts in established pathways, but rather stochastic events in individual genes that propagate through their local GRN (**Figure 3F**).

### Inter- and Intra-individual Gene Expression Stabilities are Correlated

While our analyses thus far have focused on quantifying and understanding inter-individual expression variation, intra-individual expression variation (cell-to-cell variability, or transcriptional noise) is also hypothesized to alter with age^5^. To understand the interplay between them, we calculated transcriptional noise in 207 healthy human subjects from 8 previously published single-cell RNA sequencing (scRNA-seq) datasets of the human brain^32,33^, colon^34,35^, liver^36^, lung^37^, pancreas^38^, and skeletal muscle^39^ (**Supplementary Table 5**) and compared these findings with our previous bulk-level results.

We selected only the scRNA-seq data generated using the Chromium 10x platform to ensure methodological consistency. Additionally, all data preprocessing, cleaning, clustering, and cell annotation steps were performed using a standardized pipeline applied to all studies to prevent biases introduced by differing processing protocols (**Methods**). Although several quantification methods have been proposed to measure transcriptional noise^11,12,40,41^, it remains unclear which methods are most appropriate. To be comparable with the intra-individual DVGs, we used CV to determine transcriptional noise in these samples, although a few other quantification methods are available.

Interestingly, the average transcriptional noise of a gene across all datasets shows a significant negative correlation with inter-individual stability, i.e., gGSS, across the GTEx dataset (rho: −0.294, p-value: 1.1×10^-172^; **Figure 4A**). Furthermore, the age-related increase in gGSSs positively correlates with transcriptional noise measured at the single-cell level (rho: 0.36, p-value: 1.46×10^-298^; **Figure 4B**). These results suggest that the underlying mechanisms that drive transcriptional stability operate not just between individuals, but also between cells.

We then compared transcriptional noise in young (19 y.o. < age < 40 y.o.) and old (age > 60 y.o.) samples of human brain (n = 3 young, 4 old), lung (n = 12 young, 9 old), and muscle (n = 6 young,11 old). We grouped genes into bins based on their tsGSS with 500 genes per bin. We observed distinct patterns associated with aging when examining transcriptional noise across tissues. In brain, transcriptional noise increased significantly with age across all bins, reflecting a robust and consistent age-related effect (**Figure 4C**). In contrast, lung showed no age-related increase in transcriptional noise in any bin (**Figure 4D**). Muscle exhibited an intermediate pattern, with age-related increases in transcriptional noise restricted to the bins containing the most unstable genes (**Figure 4E**). These findings highlight that the impact of age on transcriptional noise varies depending on both tissue type and gene stability, prompting us to explore these effects in greater detail within the brain, given its consistent and pronounced age-related increase in noise across all gene stability bins.

**Figure 4.**
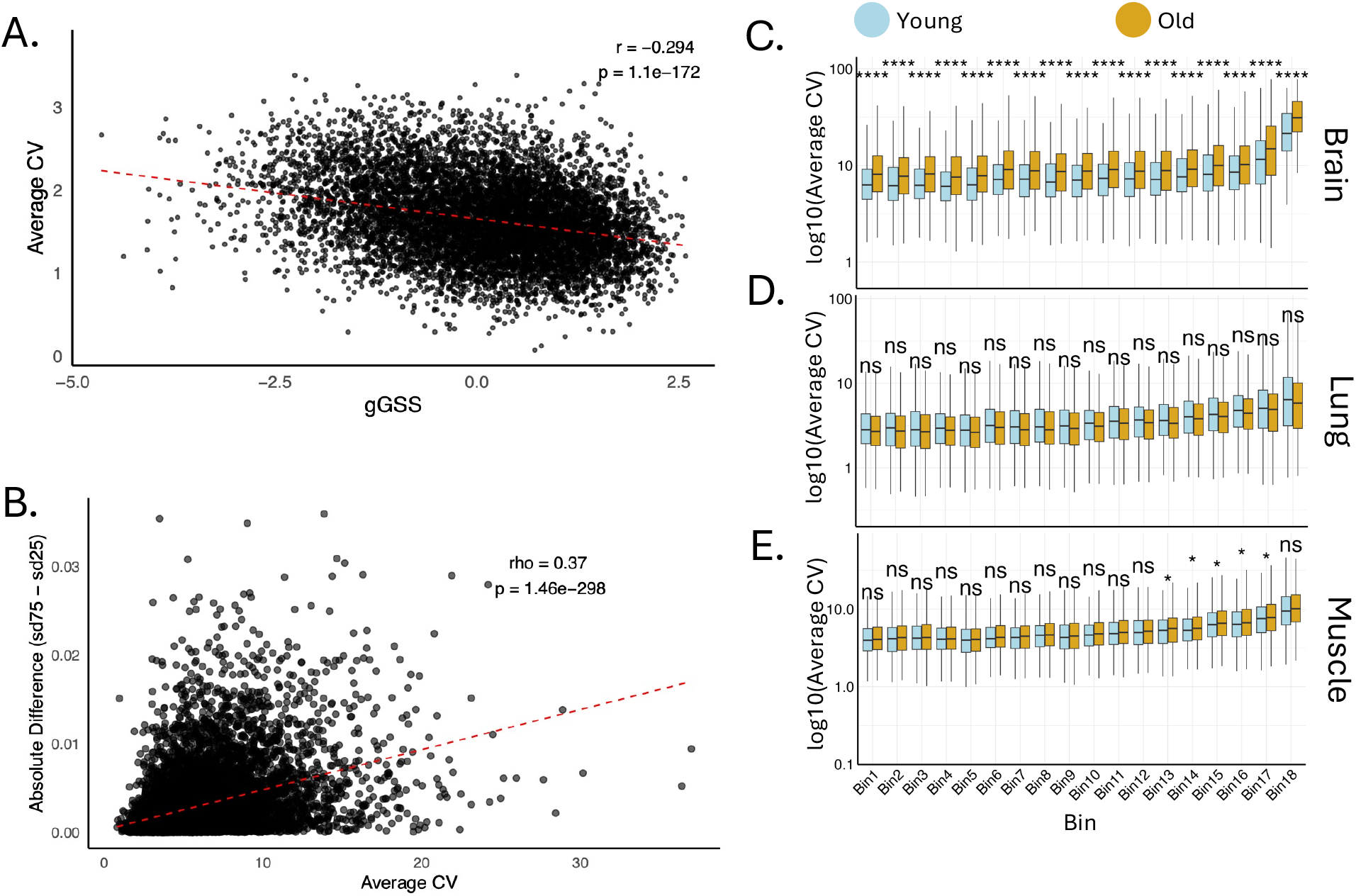
Correlation between inter- and intra-individual expression stability. (A) Correlation between gGSS calculated using bulk data and the average coefficient of variation (CV) calculated using single cell data. Each dot represents a gene, with its global gene stability score (gGSS) on the x-axis. On the y-axis, the average CV is shown, computed by first calculating the CV for that gene in each cell type within each tissue from single-cell data and then averaging across all cell types and tissues. (B) Correlation between age-related gene expression variability in bulk tissue and average CV across all cell types within each tissue from single-cell data. Age-related variability in bulk tissue was quantified for each gene as the absolute difference in standard deviation of expression between the oldest and youngest age groups. Pearson correlation rho and pvalues are reported in both A and B. Average CV for single cell data was quantified the same as in A. CV of gene expression for young (age < 40) and old (age > 60) samples in (C) brain, (D) lung, (E) muscle. Genes were grouped into bins of 500 based on their tsGSS. Bin 1 contains the 500 most stable genes, while Bin 18 contains the 500 least stable genes. Stars indicate significance from a one-tailed t-test (*p < 0.05, **p < 0.01, ***p < 0.001, ****p< 0.0001).

We focused on oligodendrocytes (n = 15,200), astrocytes (n = 3,024), and neurons (n = 5,338), as these cell types are sufficiently abundant to ensure robust sample sizes. Interestingly, the age-related increase in transcriptional noise was not uniform across these cell types: oligodendrocytes (**Figure 5A**) exhibited a significant increase with age (rho: 0.8, p-value: 0.01; **Supplementary Figure 7A**), astrocytes (**Figure 5B**) trended towards increasing noise (rho: 0.66, p-value: 0.18; **Supplementary Figure 7B**), and neurons (**Figure 5C**) showed no increase (rho: −0.25, p-value: 0.52; **Supplementary Figure 7C**). These results suggest that the impact of age on transcriptional noise within the brain is highly cell-type specific. In addition to gene level analysis, we also tested whether transcriptional noise is enriched or depleted at the network level. Notably, using GTEx data, we identified a network of interconnected, highly stable genes by selecting stable genes regulated by stable transcription factors. Centered around the *SP1* gene, we termed this subnetwork the “SP1 group”. This gene set displays remarkable inter-individual stability, suggesting that it may be intrinsically resistant to age-associated transcriptional noise. Furthermore, the SP1 group is strongly enriched for DNA damage repair pathways (**Supplementary Figure 8**), raising the possibility that DNA damage repair genes more broadly are resistant to age-associated transcriptional noise. Consistent with this hypothesis, we found that the SP1 group remained remarkably stable across all ages, even within cell types that exhibited an overall increase in noise. Similarly, DNA damage repair genes showed greater overall stability than the genome-wide background, though to a lesser extent than the SP1 group, and experienced only a modest increase in transcriptional noise with age (**Figures 5A-C**).

**Figure 5.**
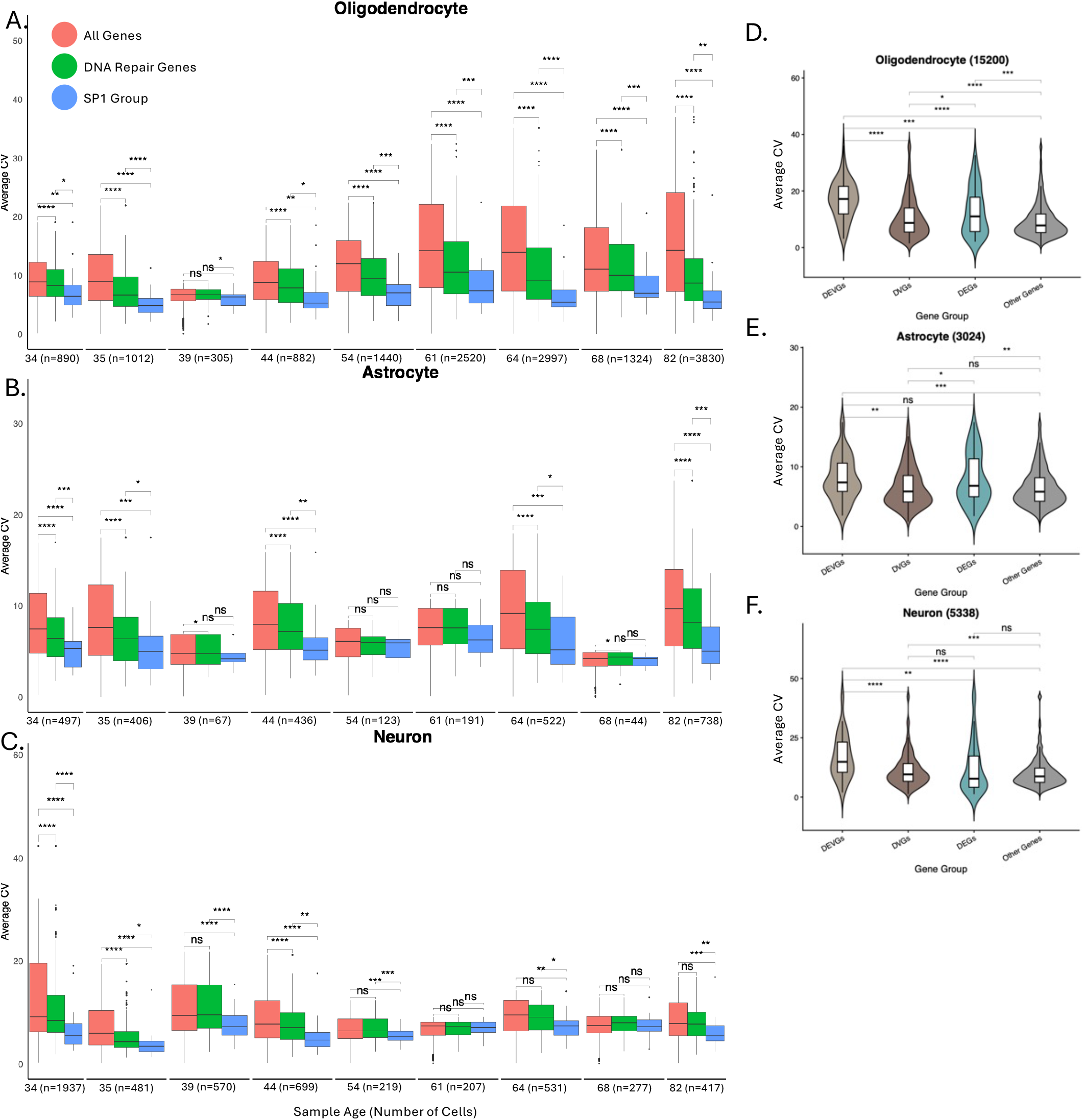
Age related increases in transcriptional noise are dependent on cell type and underlying gene regulatory network. CV of gene expression for oligodendrocytes (A), astrocytes (B), and neurons (C), within individuals. (D-F) CV of gene expression for genes grouped by bulk-level annotation. Genes were classified as DEVGs, DVGs, DEGs, of other genes based on their annotation in bulk brain data. All stars indicate significance from a two-tailed t-test (*p < 0.05, **p < 0.01, ***p < 0.001, ****p< 0.0001)

As shown above, changes in expression variation can be driven by both DVGs and DEVGs (i.e., the overlap between DVGs and DEGs; **Supplementary Table 3**) among bulk tissue samples, however it is not clear which has a more significant effect. We quantified the transcriptional noise from DEGs, DVGs, DEVGs, and the remaining genes separately for each cell type to determine the amount of variation caused by these factors (**Figure 5D-F**). Surprisingly, DEVGs produced substantially more variation on a per gene basis than DVGs, which while more numerous, had less variation per gene. Furthermore, this trend is consistent across many analyzed cell and tissue types (**Supplementary Figures 9**). This result highlights that many genes traditionally annotated as DEGs of the bulk level of changes also significantly contribute to cell-to-cell variations in gene expression.

## DISCUSSION

A recent review of the aging transcriptome highlighted nine key outstanding questions in the field, including how RNA dynamics interact across multiple hallmarks of aging^8^. Among these hallmarks, genomic and epigenomic instability are thought to influence gene expression regulation in a stochastic manner. The resulting transcriptional disruption may represent a mechanism by which these core molecular hallmarks drive higher-order phenotypes of aging, such as altered intercellular communication, inflammaging, and stem cell exhaustion. However, the underlying patterns of this transcriptional disruption remain poorly understood.

Our findings demonstrate that transcriptional alterations during aging are not limited to coordinated shifts in mean expression levels, even at the bulk tissue level. While DEGs account for the majority of observed changes, inter-individual variability, captured by DVGs, also contributes substantially, explaining approximately 7.7% of overall age-associated transcriptional alterations and up to 36% in certain pathways. These results support a model in which aging operates through two complementary modes: a shared, coordinated transcriptional program reflected by DEGs, and a heterogeneous component captured by DVGs.

One potential driver of increased inter-individual variability is heterogeneity in environmental exposures and lifestyle factors. However, such factors cannot explain the concurrent increase in cell-to-cell transcriptional variability with age, which we show affects largely the same set of genes that exhibit increased variability across individuals. Instead, these cell-to-cell differences are more likely to arise from the accumulation of stochastic genomic and epigenomic alterations that vary across cells.

This raises a key question: why do distinct sources of variation converge on the same subset of genes? Our gene regulatory network (GRN) analysis suggests that network context plays a critical role. Certain regions of the network appear to buffer transcriptional variability, whereas others may amplify it. Although the specific structural features underlying these effects remain to be defined, our results show that, compared with age-related DEGs, age-related DVGs exhibit stronger local coherence in variability and are more tightly embedded within GRN regions sharing similar variability profiles.

In addition to characterizing age-associated transcriptional variability, our analysis also provides a practical resource by identifying genes in each tissue that remain stable during aging and are suitable for use as reference genes in RT-qPCR experiments. Previous studies have demonstrated that the expression of commonly used reference genes can vary with age^28,29,42,43^. However, the identification of suitable reference genes for aging experiments remained limited, as prior studies have evaluated only a small number of candidate genes and in a few tissues. Interestingly, a recent study identified nine age-invariant genes shared across 17 tissues in C57BL/6 mice^44^; however, our results show that the human orthologs of these genes exhibit only moderate stability with age, highlighting the importance of identifying reference genes specifically suited for human studies.

In summary, our study reveals that transcriptional alterations during aging extend beyond coordinated changes in mean expression to include a substantial increase in gene-specific variability across individuals. By jointly analyzing DEGs and DVGs, we show that these two modes of transcriptional change coexist and contribute differently to the aging transcriptome. Importantly, the observation that genes exhibiting increased inter-individual variability also display elevated cell-to-cell variability suggests a shared underlying origin of transcriptional instability. Our results further indicate that this variability is not randomly distributed but is structured by gene regulatory network context, highlighting a potential role for network architecture in shaping how stochastic perturbations propagate through the transcriptome during aging.

## DATA AVAILABILITY

All gene expression datasets used in the study were published in the literature. Single cell datasets were downloaded from the Gene Expression Omnibus using accession codes: GSE126863 ^32^, GSE178341^35^, GSE192742^36^, GSE136831^37^, GSE183568^38^, GSE167186^39^, or the Human Cell Atlas^33,34^.

## ACKNOWLEDGEMENT

This work was supported by the U.S. National Institutes of Health (U19 AG056278, P01 HL160476, R35 GM159832, T32 AG029796, and U54 AG079754 and U54 AG076041). The funders had no role in study design, data collection and analysis, decision to publish, or preparation of the manuscript.

## METHODS

### Bulk RNA Sequencing Preprocessing

Raw RNA sequencing gene expression matrices were downloaded from the Genotype-Tissue Expression (GTEx) project v8 dataset. To account for differences in library size, raw counts were converted to counts-per-million (CPM) using the edgeR^45^ package. We remove lowly expressed genes that have a CPM of more than 100 in only a single sample. The 9000 genes with highest average expression across all samples were used for further analysis. Genes were identified by their Ensembl identification numbers and converted to gene names for readability. For 464 genes, multiple gene names mapped to a single Ensembl identification number; in these cases, all corresponding gene names are reported in the supplemental tables. CPM normalized counts were used to calculate GSS and DVGs. An exception to CPM normalization occurs when calculating DEGs from bulk RNA sequencing using the DESeq2^30^ pipeline, which operates directly on raw counts. Sample ages were provided as ranges rather than discrete values. To convert these into discrete values, we took the average of each range. For example, samples with an age range of 30–39 years were assigned an age of 34.5. Additionally, genes located on sex chromosomes were removed as the vast majority of their interpersonal expression variation is due to sex rather than age.

### Gene Stability Score (GSS)

GeNorm and NormFinder are tools specifically designed for analyzing qPCR data, not RNA-seq data. Although both qPCR and RNA sequencing measure gene expression levels, they differ fundamentally in how they quantify transcripts. qPCR results reflect the number of amplification cycles required to detect a transcript (Ct values), whereas RNA-seq measures the actual number of RNA molecules present by counting sequencing reads. These methodological differences make the direct application of qPCR-based normalization tools to RNA-seq data inappropriate as low qPCR results correspond to high expression while high RNA-seq results correspond to high expression. To adapt RNA-seq data for use with GeNorm and NormFinder, we transformed CPM values into pseudo-Ct values. This was done by first adding 2 to all CPM values to avoid issues with zeros, then applying a log2 transformation followed by inversion (**Equation 2**).

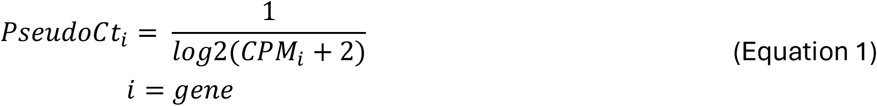

This process converts high CPM values (high expression) into low pseudo-Ct scores (also representing high expression), maintaining the interpretative consistency with qPCR data. Adding 2 before transformation ensures that genes with zero CPM receive appropriately high pseudo-Ct scores, reflecting low expression. We then subsetted this dataset down to the top 9000 genes with the highest average expression across all samples. We retained only the 9000 most highly expressed genes because GeNorm and NormFinder are designed to evaluate candidate benchmark genes with strong expression. Their suitability for analyzing lowly expressed genes, which are more susceptible to technical artifacts such as dropouts, remains uncertain. Finally, we applied GeNorm and NormFinder to these pseudo-Ct scores to assess gene stability. Additionally, we calculated the Coefficient of Variation (CV) on the pseudo-Ct data. Principal Component Analysis (PCA) was performed separately for each tissue and across all tissues combined (**Supplementary Figure 4**). In all cases, the first principal component captured the vast majority of the variance (**Supplementary Figure 2**), supporting its use as a robust quantitative measure of gene stability. We refer to this metric as the Gene Stability Score (GSS).

### Gene Regulatory Network Analysis

Gene Regulatory Network (GRN) data was downloaded from the Transcriptional Regulatory Relationships Uncovered by Sentence-base Text mining (TRRUST) v1 database^31^. This database uses a sentence-based text mining algorithm to scan the preexisting literature and pull out previously described regulatory relationships. At the time of publication the TRRUST website is not operational, however, gene regulatory relationships can still be found at: https://github.com/slowkow/tftargets/blob/master/data-raw/TRRUST/trrust_rawdata.txt.gz. To generate the stability similarity metric, we first ranked all genes based on their previously defined GSS. For each gene, we then computed a raw similarity score by calculating the average absolute difference in stability between the gene and its immediate neighbors within the GRN (**Equation 2**). To assess whether the observed stability similarity was greater or less than expected by random chance, we employed a bootstrapping approach. For each gene, we randomly sampled the same number of genes as its actual number of neighbors in the GRN and calculated the average stability difference between the gene and these randomly selected genes. This random sampling procedure was repeated 100 times, and the resulting values were averaged to produce a bootstrapped similarity score, representing the level of stability similarity a given gene would be expected to have with its local GRN under random chance (**Equation 3**). Finally, we defined the stability similarity score as the log fold change between the bootstrapped and raw similarity scores (**Equation 4)**.

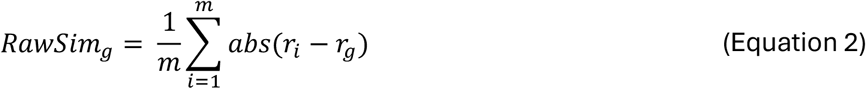

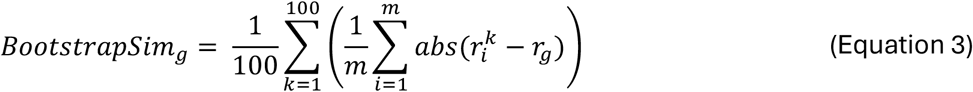

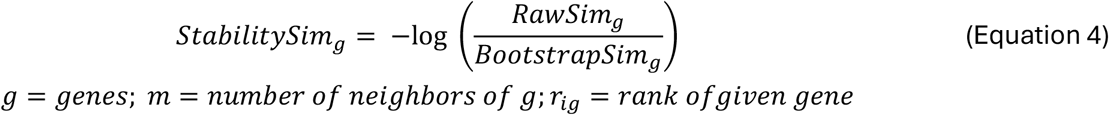

### Quantifying DEGs and DVGs

DEGs from bulk RNA sequencing were quantified using the DESeq2^30^ pipeline with age modeled as a continuous covariate. Genes were labeled as DEGs if they had a yearly absolute Log2FC of greater than 0.0116 and an adjusted p-value less than 0.05. This cutoff represents a doubling or halving of a gene’s expression over 60 years, and is used when modeling age as a continuous covariate. To quantify DVGs we compared the standard deviation of gene expression between individuals within defined groups. Age-related DVGs were calculated by comparing individuals in the youngest age bracket with those in the oldest age bracket. Sex-related DVGs were calculated by comparing male and female groups. To test whether the observed differences in variance were statistically significant, we applied Levene’s test and labeled genes as DVGs if it had an adjusted p-value less than 0.05.

While Levene’s test identifies whether variability differs between age extremes, it does not indicate whether variability systematically changes with age. To address this, we calculated the SD of expression for each gene within all age groups, then ranked the age groups by increasing variance. By testing for correlation between age and variance using Spearman Correlation, we were able to assess whether gene expression variability consistently increases or decreases with age, suggesting a progressive, age-related change in variability.

### Measuring Contributions to GSS

To measure the relative contributions of DEGs and DVGs arising from age and sex separately we modeled our previously calculate GSS score as a linear combination of these factors. While other factors, like environmental, certainly affect gene stability, we are unable to model these factors due to limitations in the metadata provided by GTEx. We used the relaimpo^46^ R package to compute the relative importance of the factors in our model. This package decomposes the models *R*^*2*^ into contributions from each predictor, as well as the variance that is unexplained by the model. Only genes that met our DEG or DVG significance thresholds (padj < 0.05) across all four predictors (age DEGs, age DVGs, sex DEGs, and sex DVGs) were included in the regression. This filtering ensures that the contributions measured for each predictor reflect robust, reproducible associations with age- or sex-related changes in expression level or variability.

### Single Cell RNA Sequencing Pipeline

All scRNA-seq datasets were processed using a standardized pipeline we previously developed. Briefly, this pipeline takes raw expression matrices and performs a series of steps including data cleaning and quality control, normalization, integration, clustering, and cell type annotation. During quality control, we remove outlier cells that are either poorly sequenced or identified as potential doublets. Normalization is then applied within each study to correct for differences in library size. Integration is used to mitigate technical variability between samples. The integrated data is subsequently clustered, and each cluster is annotated based on canonical marker gene expression. Using the resulting cleaned and annotated datasets, we perform differential gene expression analysis within each cluster using Nebula^47^ and quantify transcriptional variability using the Coefficient of Variation.

