## Supplemental Figures 1-9 for "Age-associated increases in inter-individual gene expression variability across human tissues"

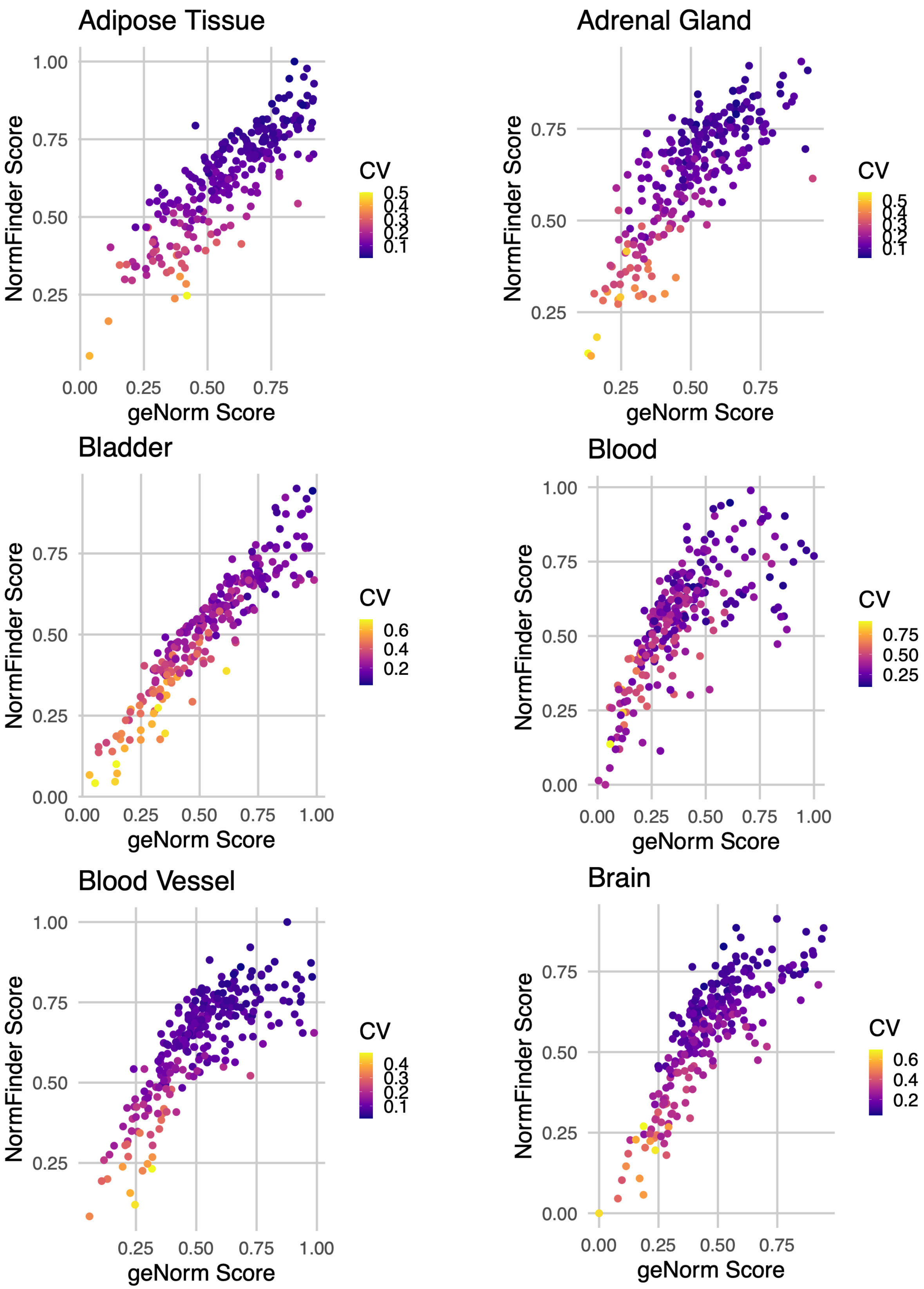


**Figure S1. Comparison of Normfinder, geNorm, and CV for genes in each tissue.** Normfinder and geNorm scores are normalized and data from each tissue is downsampled to 250 genes for readability.

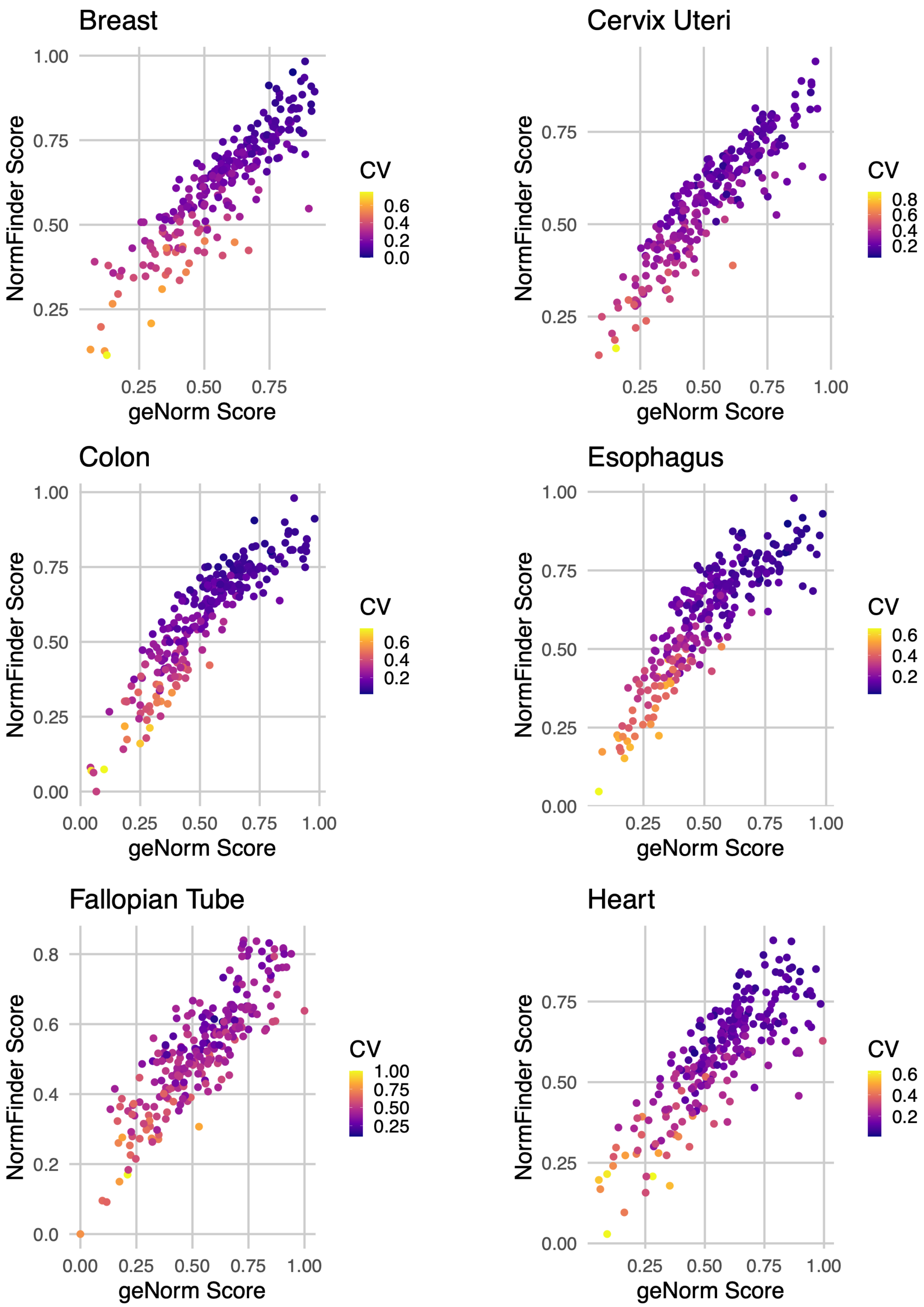


**Figure S1 continued.**

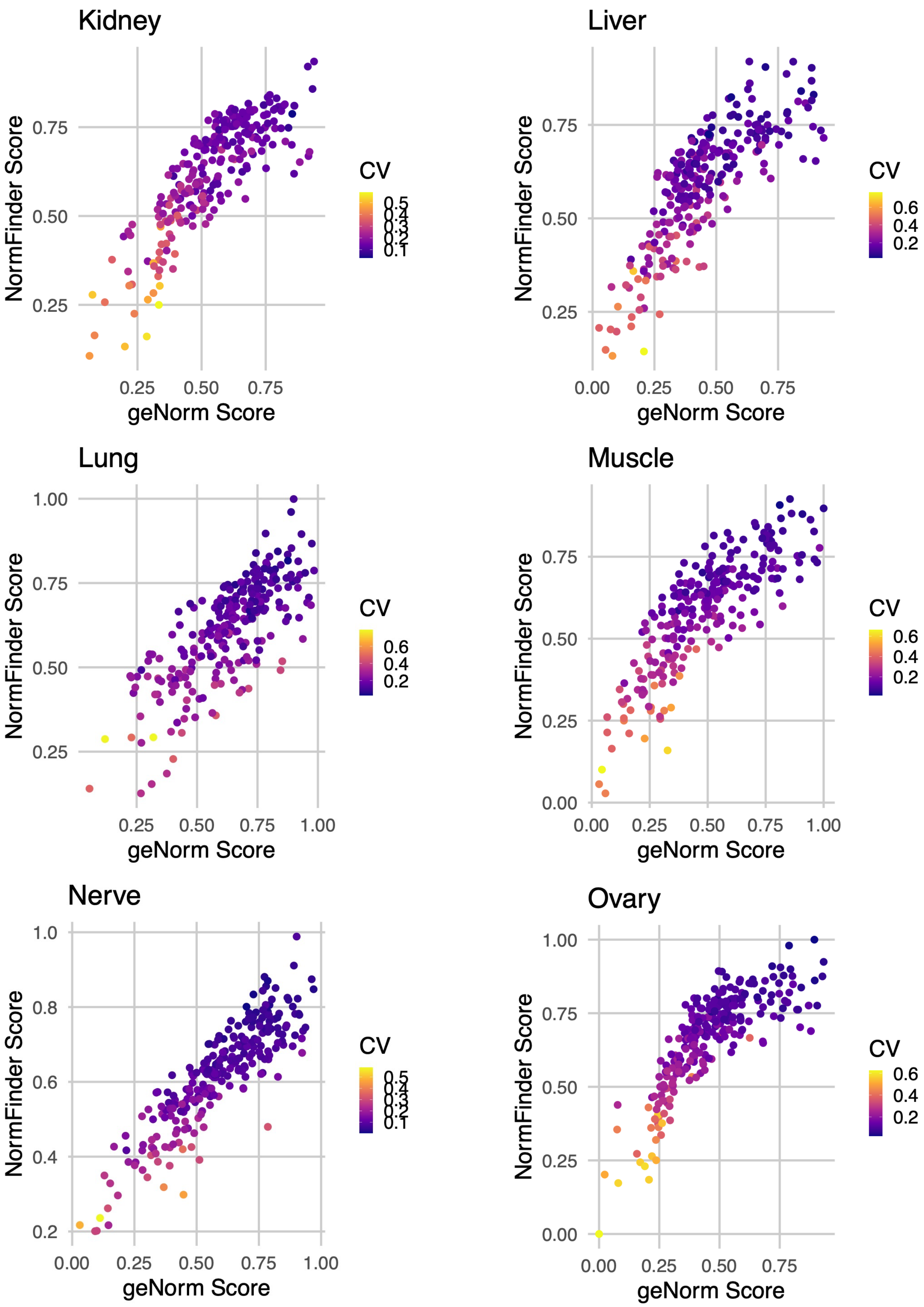


**Figure S1 continued.**

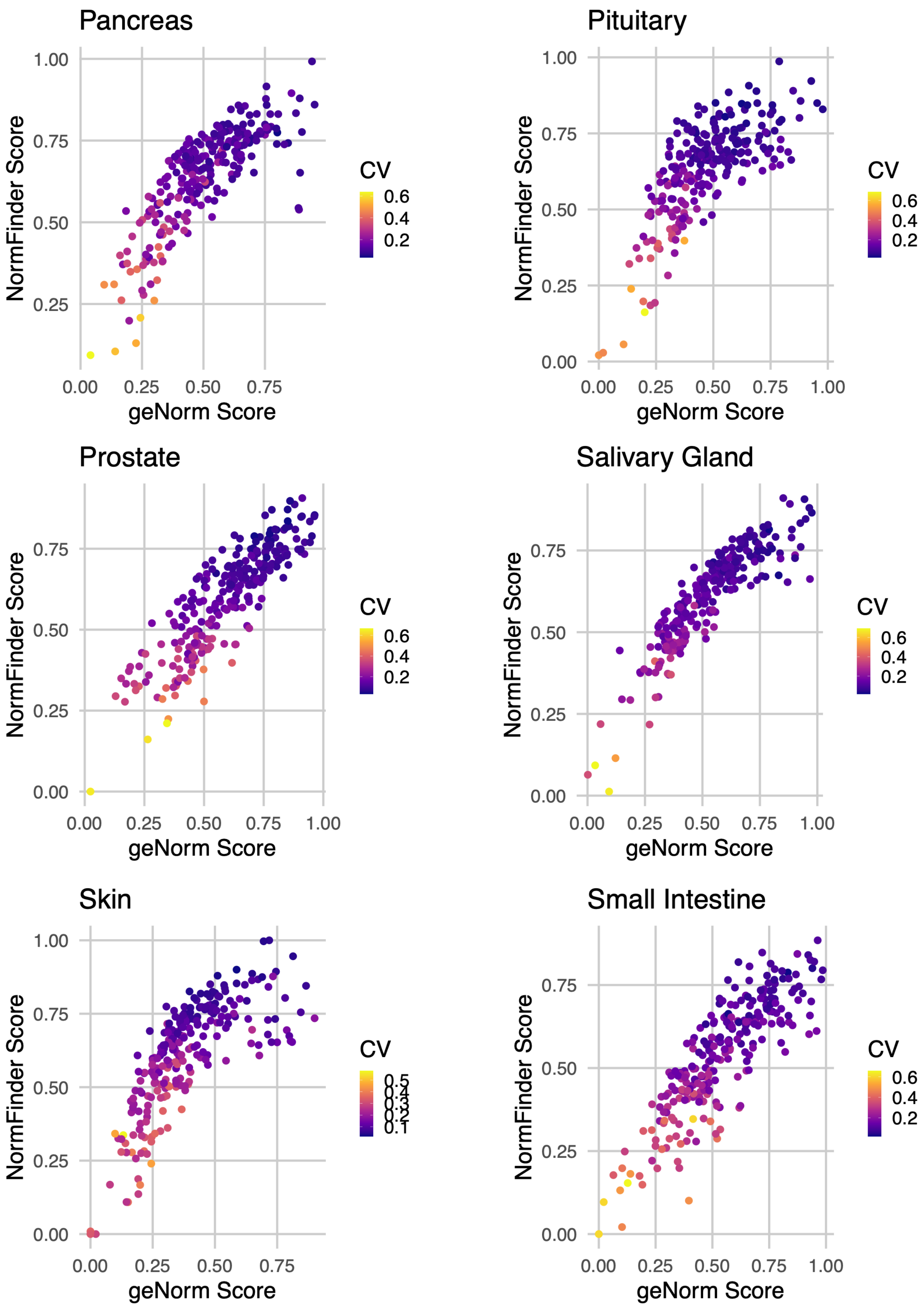


**Figure S1 continued.**

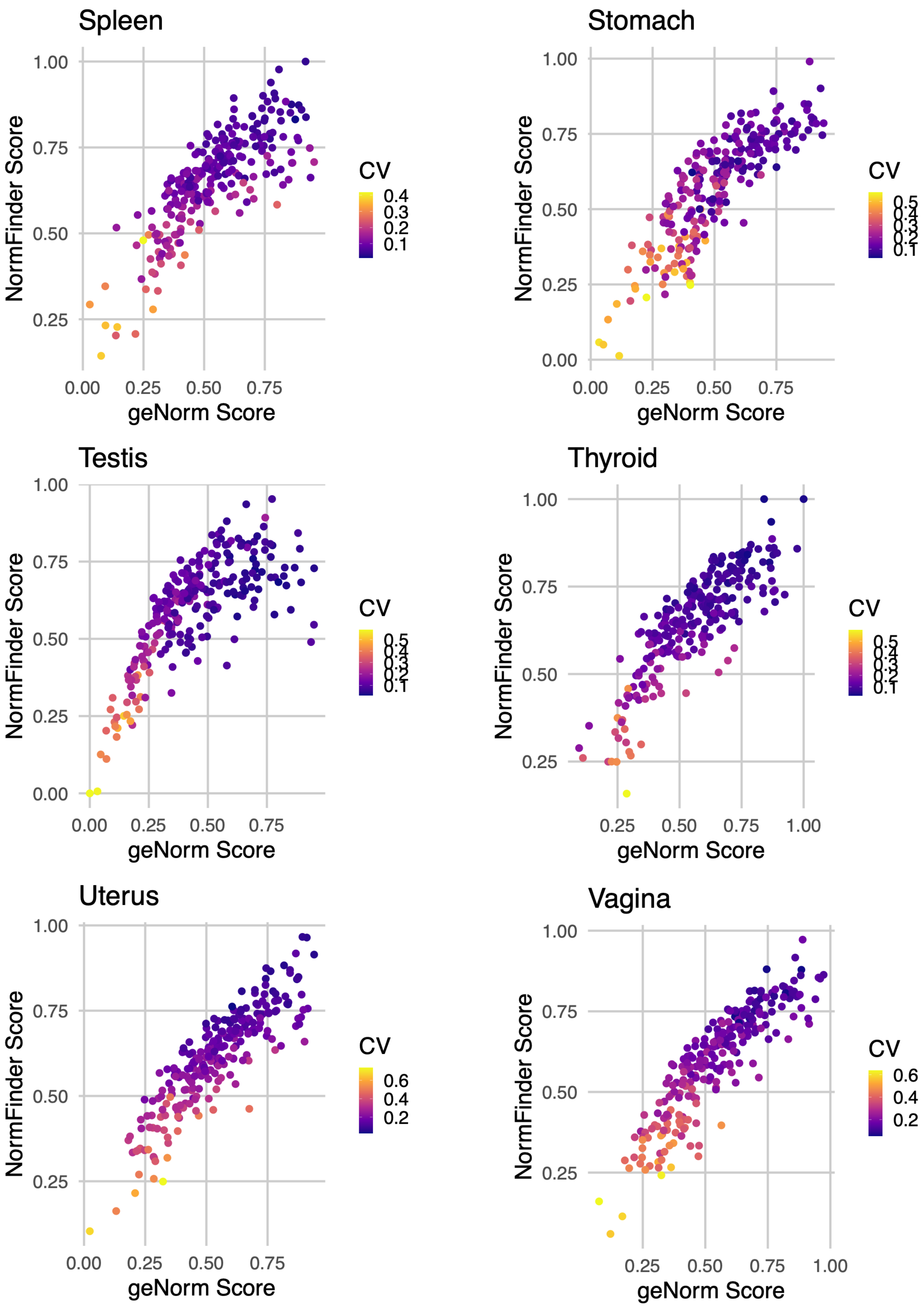


**Figure S1 continued.**

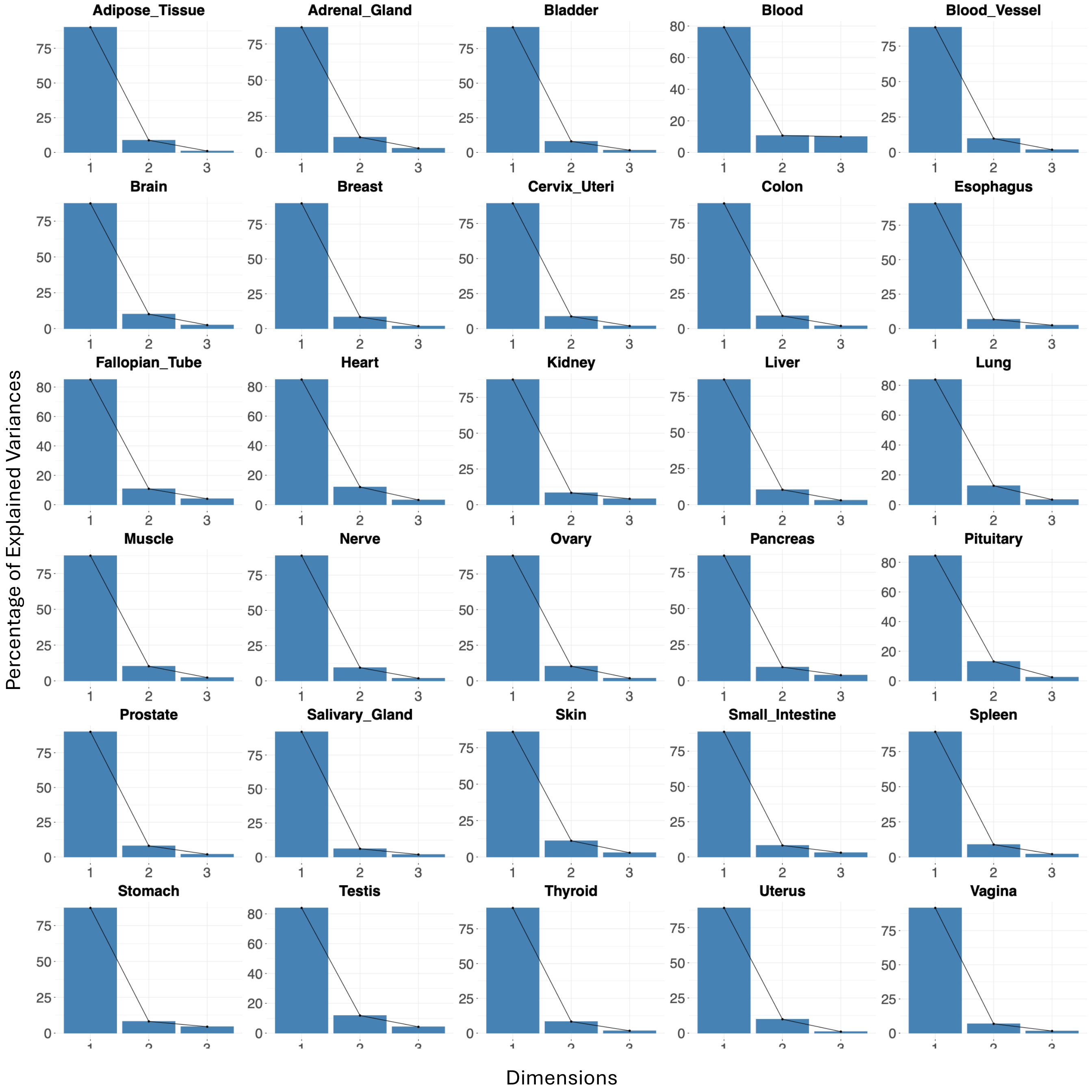


**Figure S2. Scree plots for each tissue.** PCA reductions were performed on Normfinder, geNorm, and CV scores for each tissue. In all tissues, the first principal component explains the vast majority of variance.

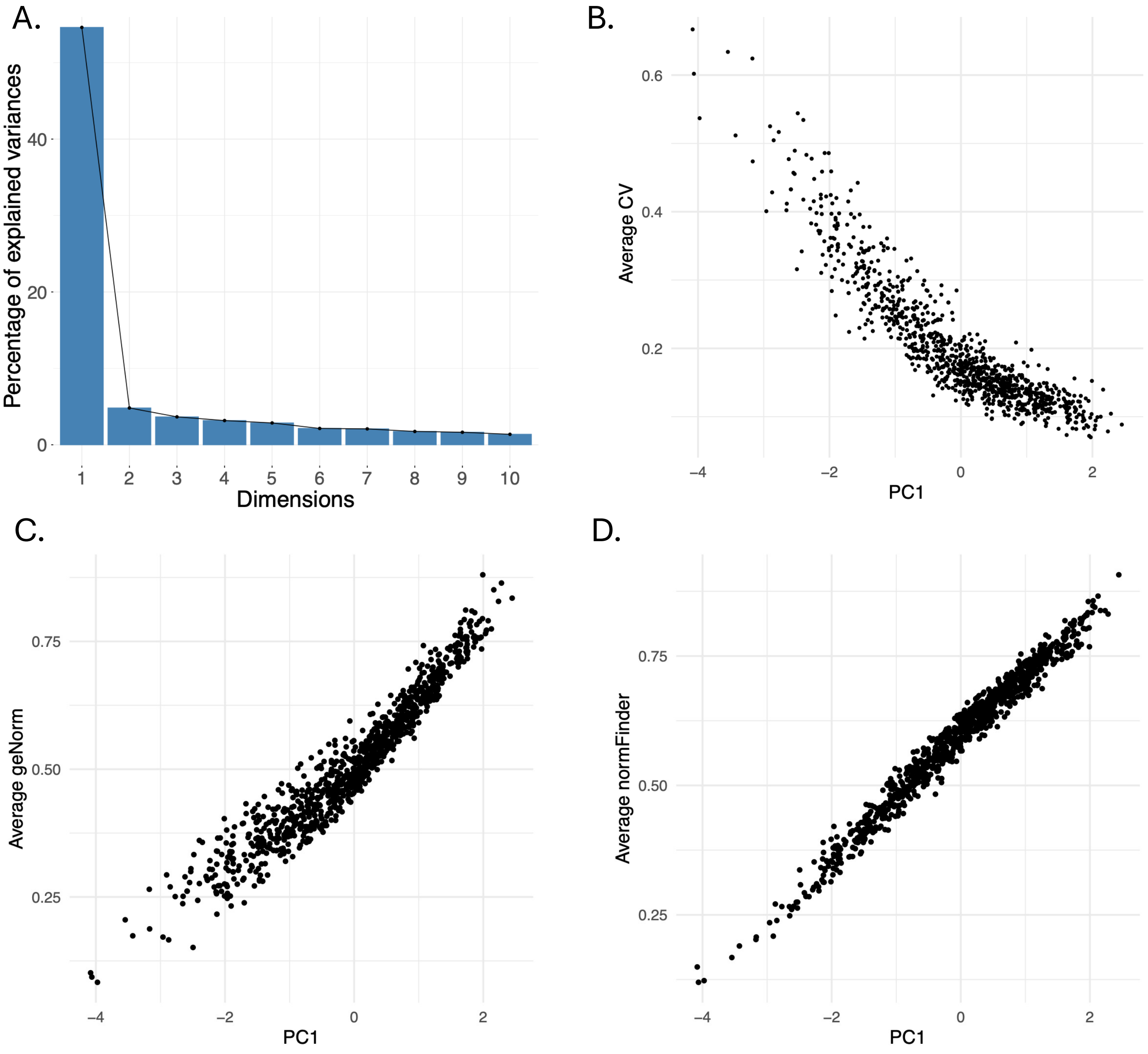


**Figure S3. Validation of global PCA reduction.** PCA reduction was performed using the NormFinder, geNorm, and CV scroes from all 30 tissues. (A) Scree plot of the first 10 principal components shows that the vast majority of variance can be explained by the first principal component (PC1). Correlation of PC1 with (B) average CV across all tissues, (C) average geNorm across all tissues, and (D) average NormFinder across all tissues.

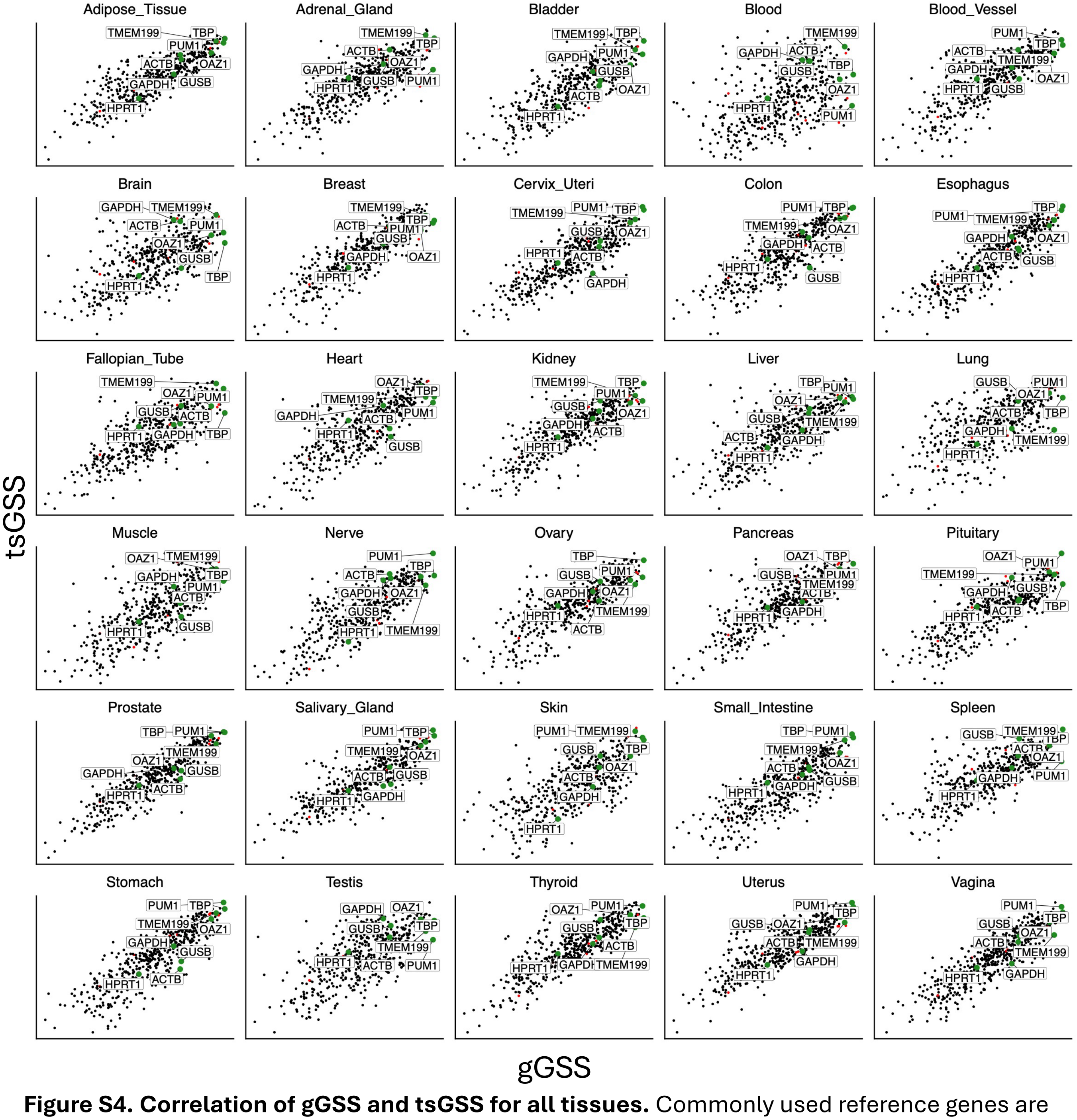


**Figure S4. Correlation of gGSS and tsGSS for all tissues.** Commonly used reference genes are labeled and data from each tissue is downsampled to 1000 genes for readability

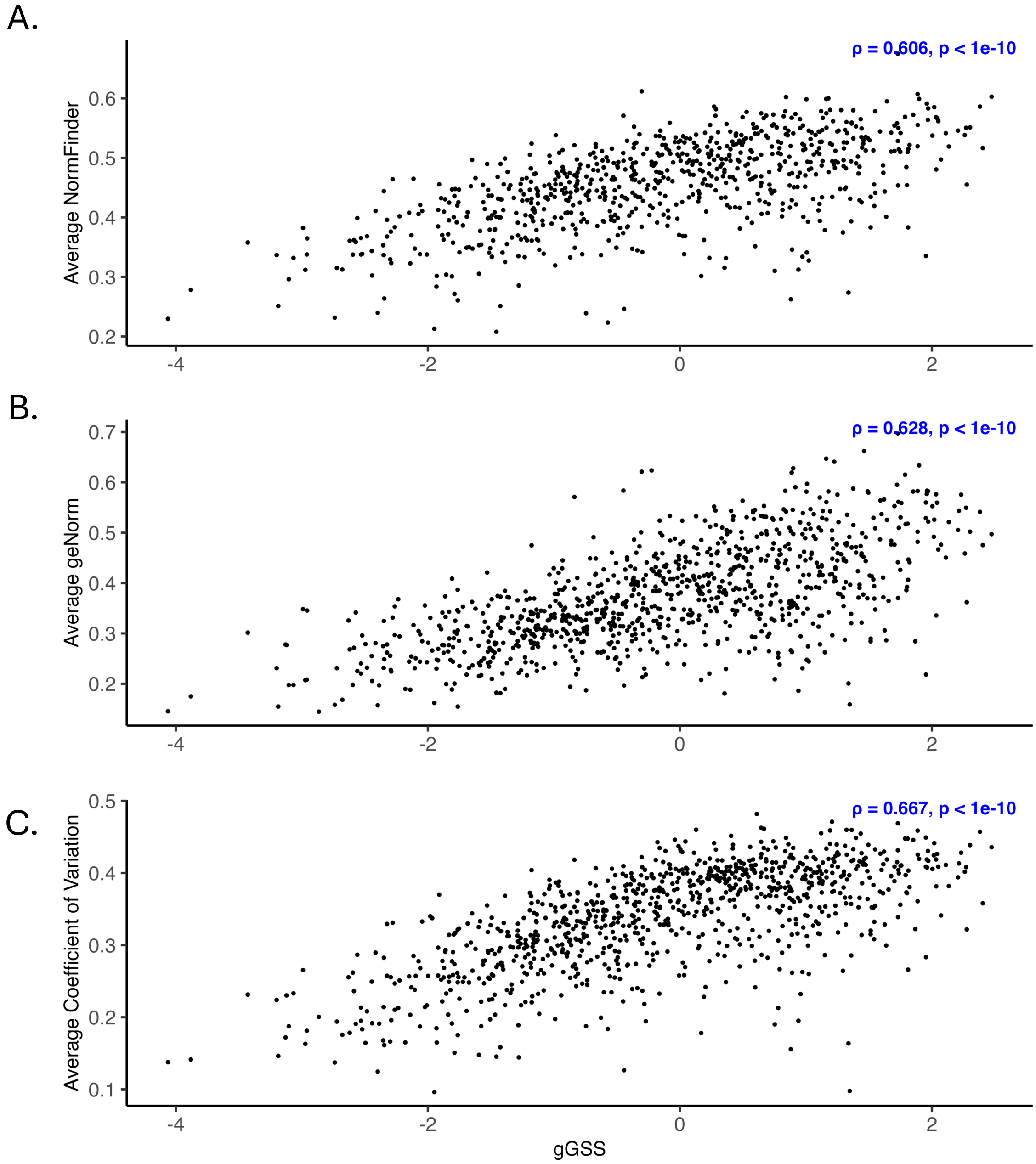


**Figure S5. Correlation of stability scores derived from GTEx and Human Protein Atlas.** The gGSS derived from the GTEx dataset was compared to (A) the average NormFinder score, (B) the average geNorm score, and (C) the average CV derived from the Human Protein Atlas.

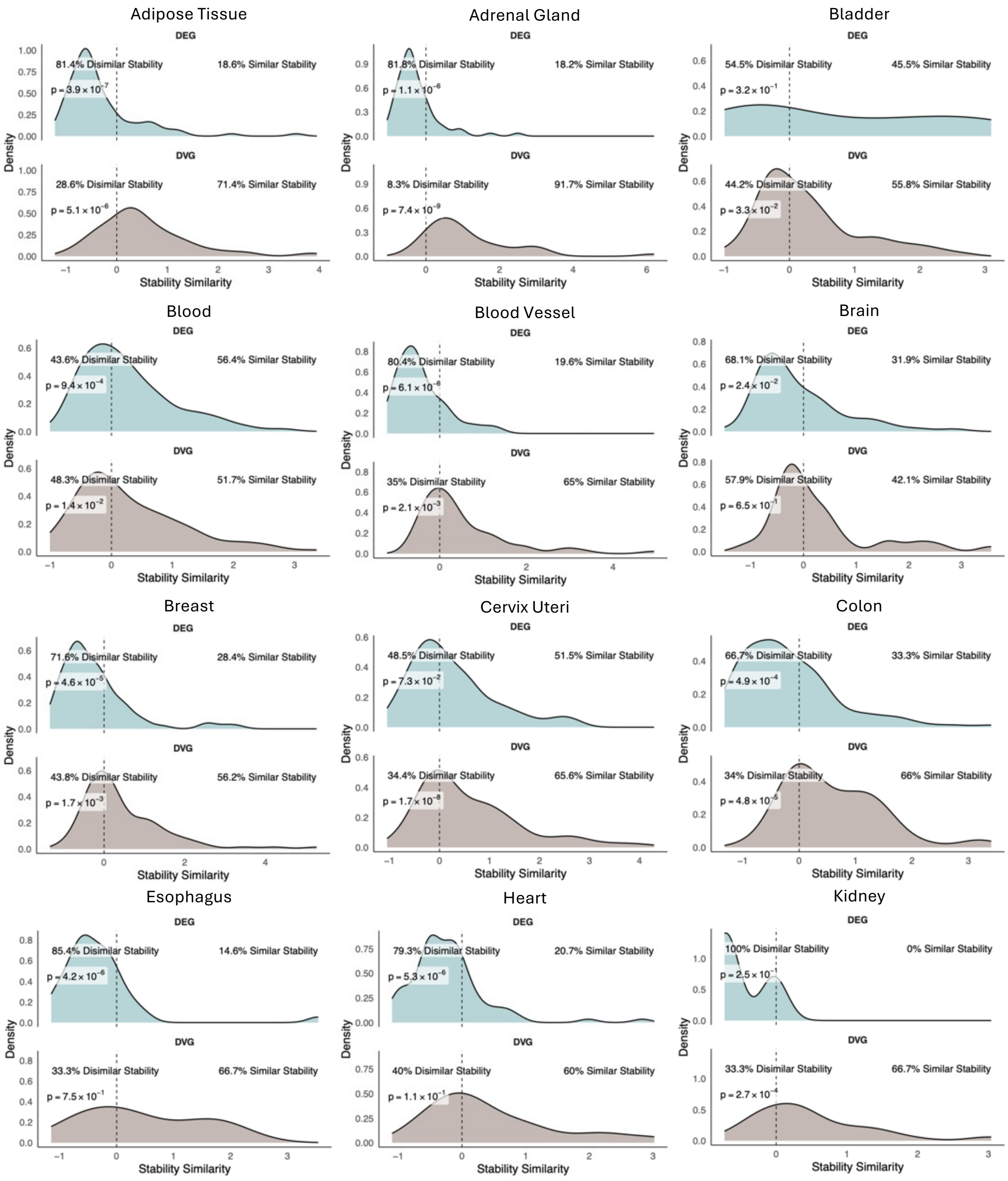


**Figure S6. Density plots of stability similarity scores for DEGs and DVGs in each tissue.** Displayed p-values are from a two-sided Wilcox test assessing whether the observed stability similarity distribution differs from 0.

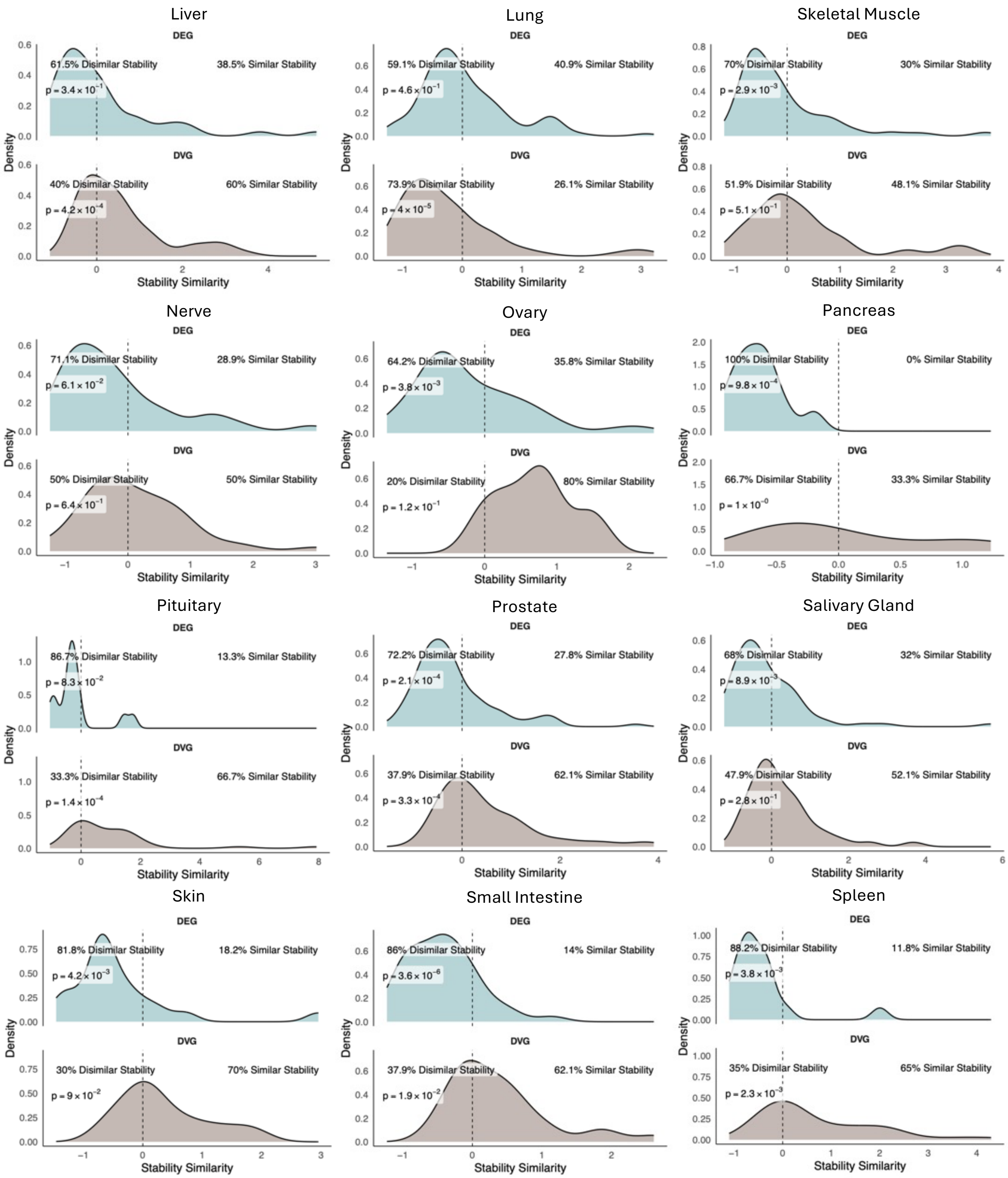


**Figure S6 continued**

Stomach

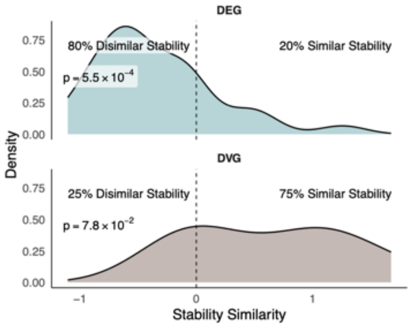


**Figure S6 continued**

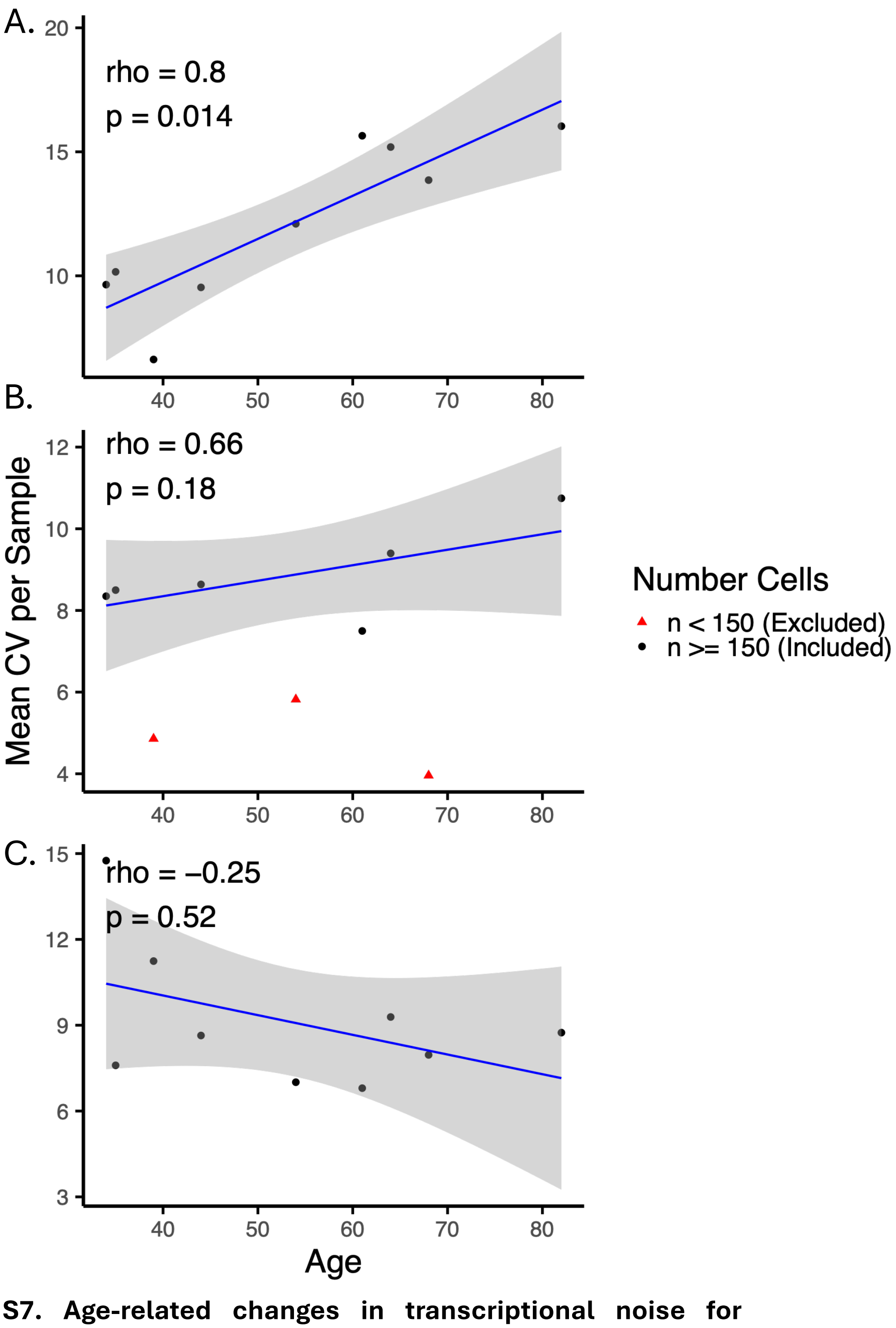


**Figure S7. Age-related changes in transcriptional noise for selected brain cell types.** The average CV for all genes in (A) Oligodendrocytes, (B) Astrocytes, and (C) Neurons for each sample are displayed. Displayed rho and p-values are from a Spearman correlation between average CV and age. Samples with fewer than 150 cells are shown in red but were excluded from correlation analysis.

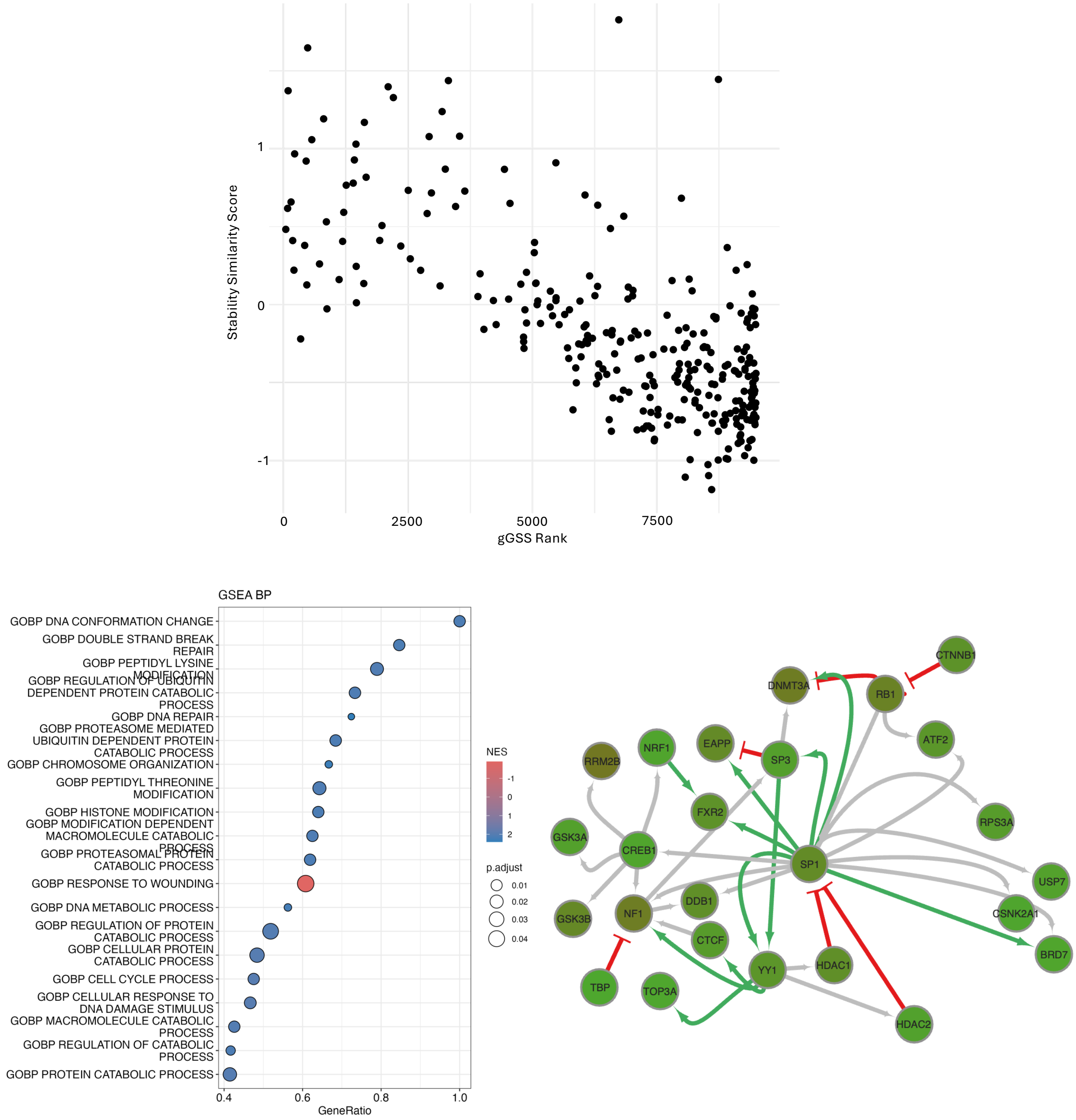


**Figure S8. The SP1 group.** (A) Scatterplot comparing the stability similarity of a gene with its gGSS ranking. High stability similarity score indicates a genes is more similar in stability to its neighbors than is expected by random change, while low scores indicate that it is less similar than expected by random change. Low gGSS rank indicates stability while high gGSS rank indicates instability. Because these metrics are correlated, we performed a PCA reduction to to reduce dimensionality and capture the main axis of variation. Genes were then ranked along the primary PCA component and (B) GSEA enrichment analysis was performed using this ranking. (C) The SP1 group's regulatory network is displaywed where green arrows represent activation, red "blockers" represent suppression, and grey arrows represent interactions of unknown direction.

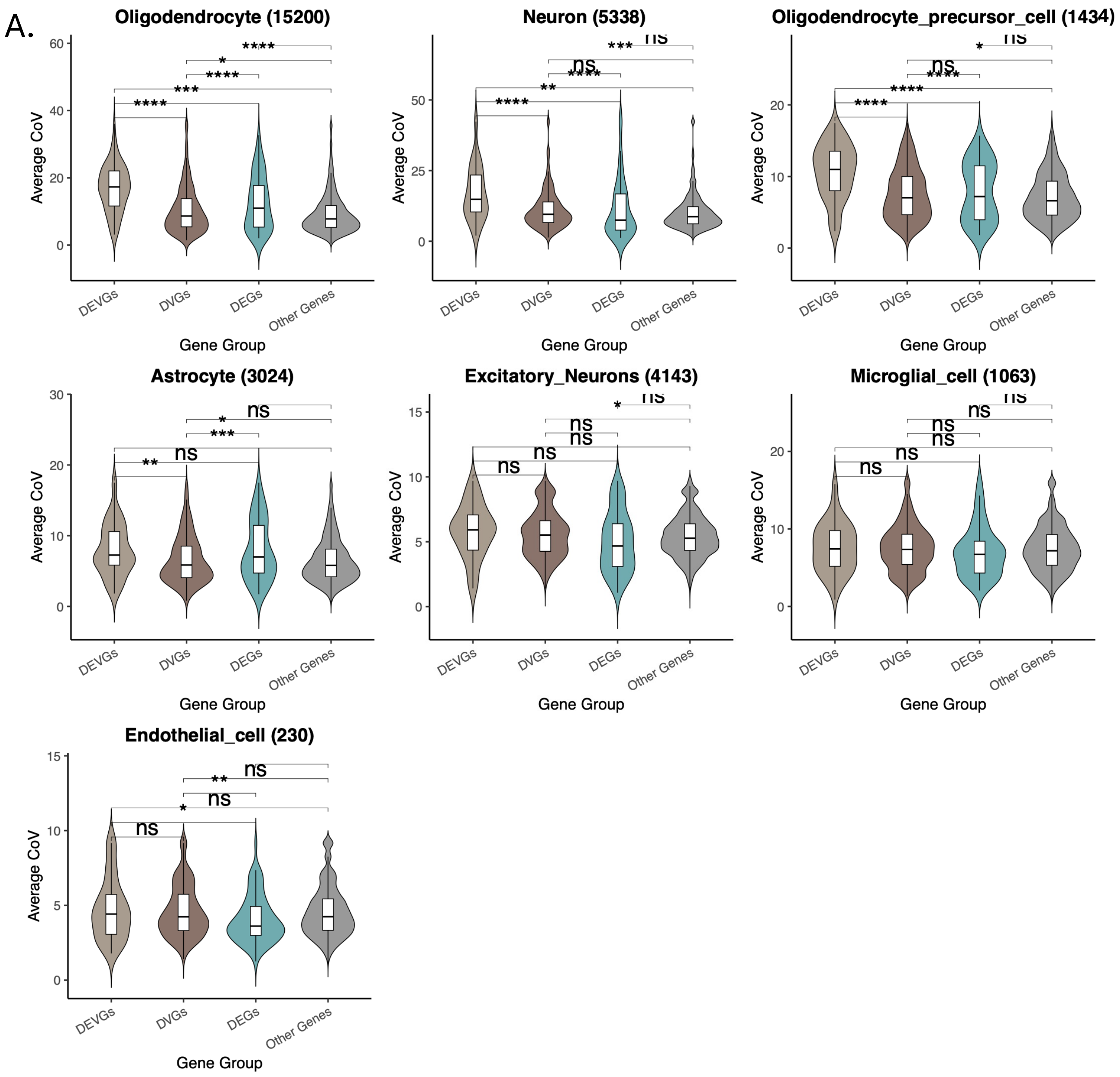


**Figure S9. Transcriptional Noise of DEVGs, DVGs, and DEGs.** Average coefficient of variation of genes based on bulk level annotations for cell types from (A) brain, (B) colon, (C) pancreas, and (D) muscle.

B.

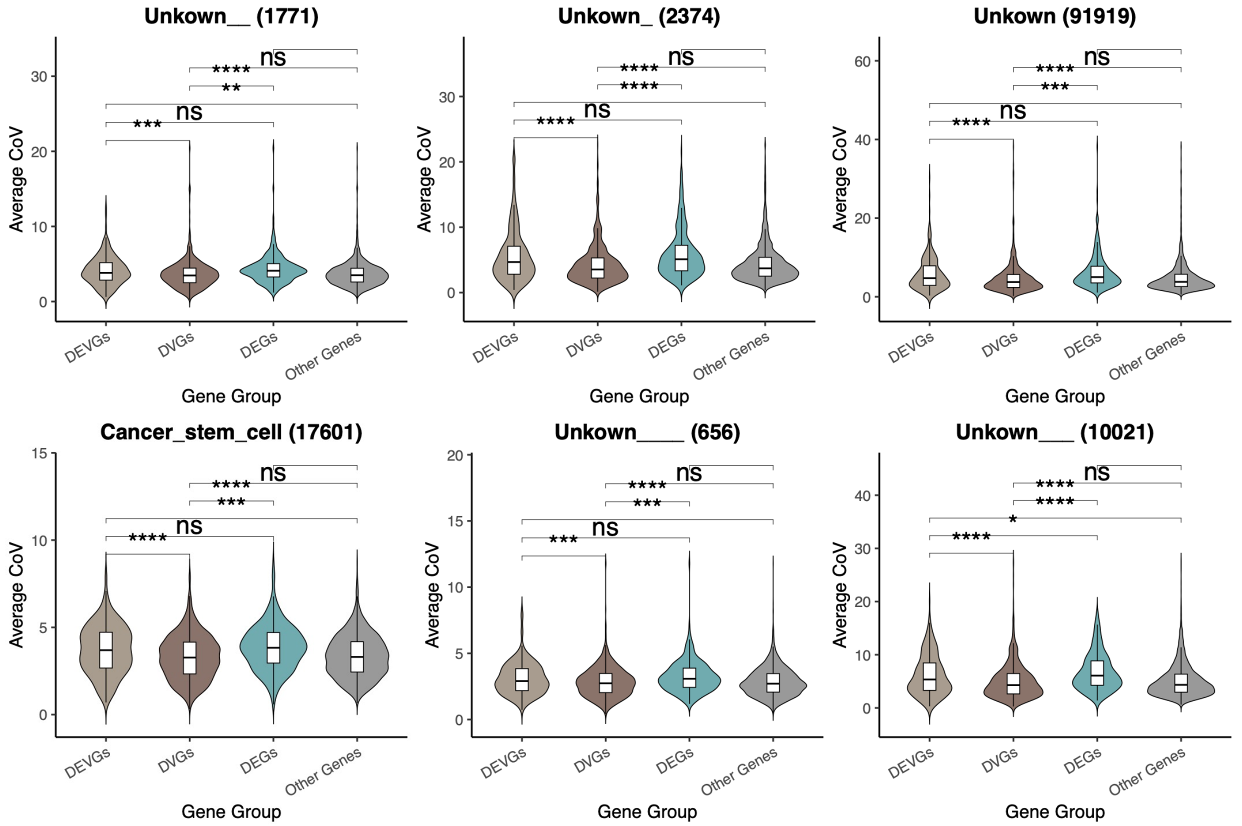


C.

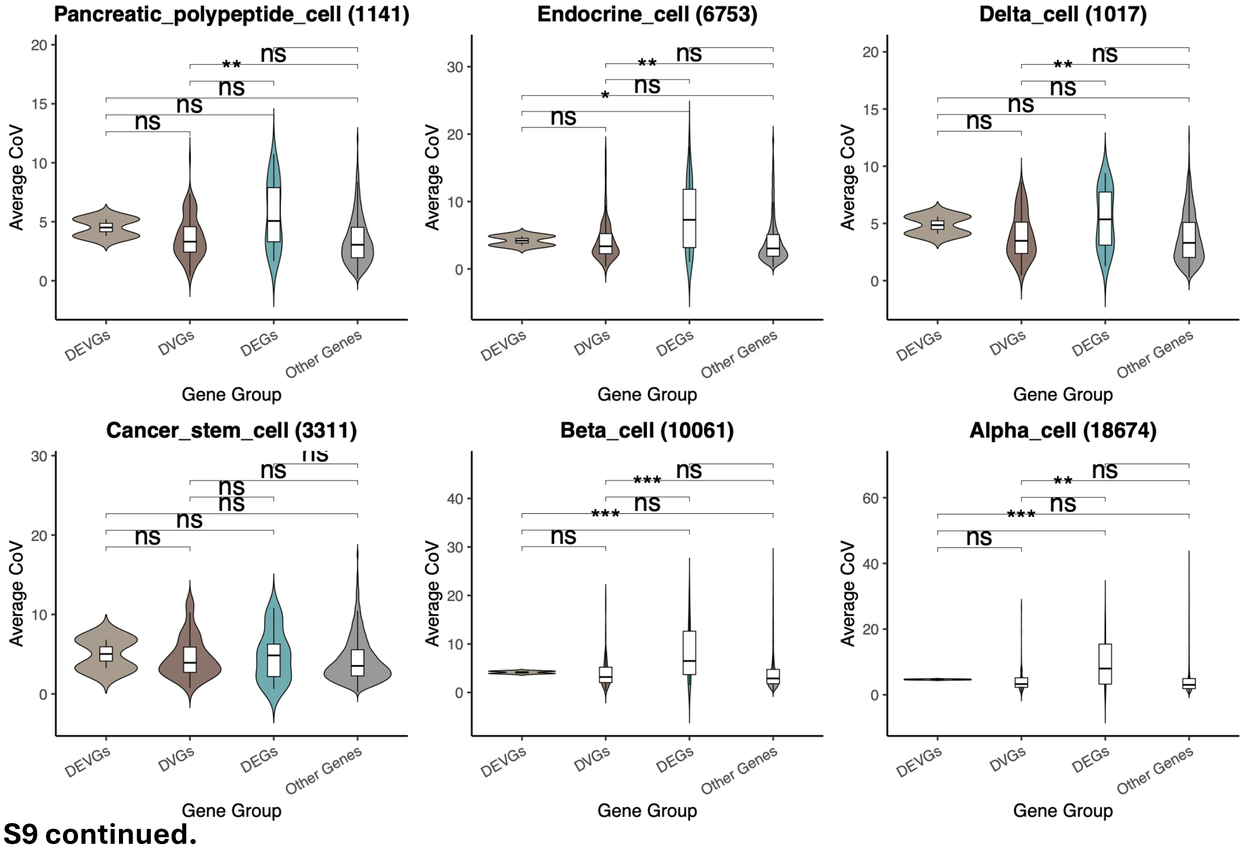


**Figure S9 continued.**

D.

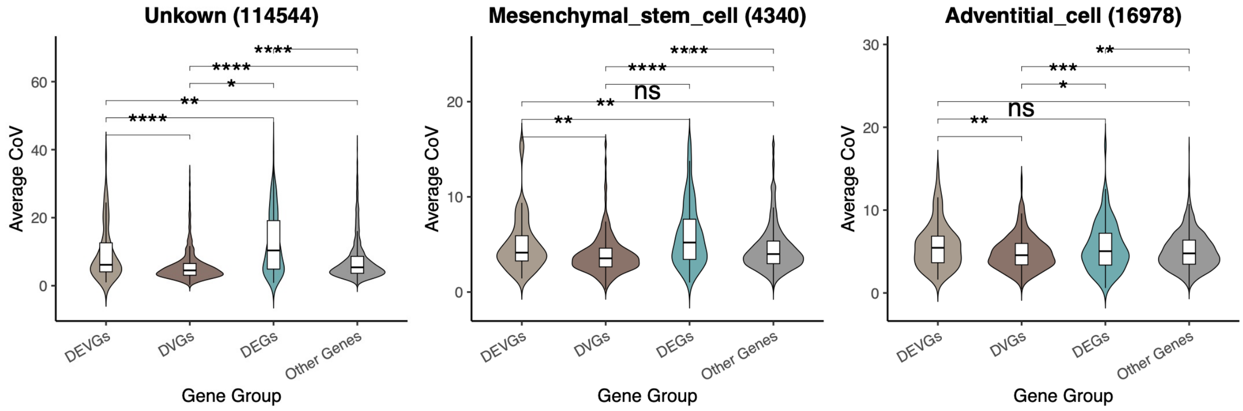


**Figure S9 continued.**
